# A Population Genomics Analysis of the Native Irish Galway Sheep Breed

**DOI:** 10.1101/645754

**Authors:** Gillian P. McHugo, Sam Browett, Imtiaz A. S. Randhawa, Dawn J. Howard, Michael P. Mullen, Ian W. Richardson, Stephen D. E. Park, David A. Magee, Erik Scraggs, Michael Dover, Carolina N. Correia, James P. Hanrahan, David E. MacHugh

## Abstract

The Galway sheep population is the only native Irish sheep breed and represents an important livestock genetic resource, which is currently categorised as “at-risk”. In the present study, comparative population genomics analyses of Galway sheep and other sheep populations of European origin were used to investigate the microevolution and recent genetic history of the breed. These analyses support the hypothesis that British Leicester sheep were used in the formation of the Galway breed and suggest more recent gene flow from the Suffolk sheep breed. When compared to conventional and endangered breeds, the Galway breed was intermediate in effective population size, genomic inbreeding and runs of homozygosity. This indicates that, although the Galway breed is declining, it is still relatively genetically diverse and that conservation and management plans informed by genomic information may aid its recovery. The Galway breed also exhibited distinct genomic signatures of artificial or natural selection when compared to other breeds, which highlighted candidate genes that may be involved in meat and wool production.

## INTRODUCTION

Sheep were domesticated more than 10,000 years ago and have since been bred for a variety of uses including meat, milk and wool production (Taberlet et al., 2011; Larson and Fuller, 2014; MacHugh et al., 2017). During the last 50 years, the focus of the global sheep industry on only a subset of the 1,400 recorded sheep breeds with enhanced productivity and high-quality outputs has resulted in many locally-adapted (local) breeds becoming endangered or extinct (Taberlet et al., 2008; Kijas et al., 2009; Kijas et al., 2012). These breeds are generally considered independent genetic units because crosses are usually not used for further reproduction (Taberlet et al., 2008). Local or heritage livestock breeds are important because they constitute reservoirs of biological diversity which may be important genetic resources for domestic animal species in the face of climate change and increased food requirements in the future (Taberlet et al., 2008; Bowles, 2015). In particular, functionally important natural sequence variants (NSVs) identified in the genomes of local or heritage breeds may become increasingly important as targeted genome editing technologies are employed in genetic improvement programmes (Wells, 2013; Petersen, 2017; Van Eenennaam, 2017).

The local sheep breeds on the periphery of Northern Europe are recognised as heritage livestock populations that should be conserved and represent important sources of novel genetic diversity accumulated over centuries of microevolution and adaptation to marginal agroecological environments (Tapio et al., 2005). In this regard, the Galway sheep breed is the only surviving sheep breed native to Ireland (Curran, 2010); it was once the principal lowland sheep breed in the west of the country, but is now considered at-risk by the Food and Agriculture Organization (Food and Agriculture Organization, 2019).

The Galway breed is thought to have originated in the 1840s and is likely to have emerged as a composite breed from several indigenous and imported sheep populations present in Ireland at that time (Hanrahan, 1999), including the important Dishley or New Leicester foundational breed developed by Robert Bakewell (Wykes, 2004). However, it was not until 1923 that a formal herd book was established (Curran, 2010; Food and Agriculture Organization, 2019). Therefore, the range of sheep populations ancestral to the Galway breed in the 18^th^ and 19^th^ centuries, coupled with the possibility of more recent gene flow poses questions concerning the genetic distinctiveness and admixture history of the breed. In addition, the Galway breed has declined from a peak population size in the 1960s when it was the focus of lowland sheep farming in western Ireland (Martin, 1975a; Raftice, 2001; Curran, 2010). By 1994, as defined by the UK Rare Breeds Survival Trust, the Galway breed had reached “critical” status for sheep breeds with only 300 pedigree breeding ewes registered (Curran, 2010). Since being classed as endangered by the Irish Government in 1998, the number of pedigree Galway sheep has increased due to conservation efforts; however, the breed population size is currently decreasing, raising concerns regarding remaining genetic diversity and the overall viability of the population (Curran, 2010; Food and Agriculture Organization, 2019).

As a local breed with a low census population size, the main threat to the long-term survival of the Galway breed is replacement by more productive commercial breeds, which would further reduce the population size, reduce genetic diversity and increase inbreeding. Other challenges faced by threatened local livestock breeds include increased genetic drift, poor animal husbandry and management, deliberate or inadvertent crossbreeding and geographical isolation, which increases the risk of extinction (Taberlet et al., 2008; Allendorf et al., 2013). In recent years, with the availability of increasingly powerful genomics technologies, a conservation programme for Galway sheep has been proposed that would leverage molecular genetic information (McHugh et al., 2014). McHugh and colleagues also propose that genome-enabled breeding (genomic selection) could be used in threatened livestock populations to improve production, health and reproduction traits, thereby decelerating replacement by modern breeds (Biscarini et al., 2015).

To provide information that may be relevant to genetic conservation of the Galway sheep breed, in the present study we performed high-resolution population genomics analyses in conjunction with 21 comparator breeds of European origin. These analyses included multivariate analyses of genomic diversity, phylogenetic network graph reconstruction, evaluation of genetic structure and inbreeding, modelling of historical effective population sizes and functional analyses of artificial and natural selection across the Galway sheep genome.

## MATERIALS AND METHODS

### Galway and Irish Suffolk Sheep DNA Sampling

The Galway and Irish Suffolk sheep DNA samples used for the current survey were generated from peripheral blood samples collected in standard heparinised Vacutainer blood collection tubes (Becton-Dickinson Ltd., Dublin, Ireland). High-quality genomic DNA was then purified from 200 µl of blood from each animal using standard laboratory methods (Howard, 2008).

### Additional SNP Data Sources and Data Filtering

High-density SNP data were obtained from the International Sheep Genomics Consortium Sheep HapMap Project and consisted of 2,819 sheep from 74 breeds genotyped for 49,034 evenly-spaced SNPs using the Illumina^®^ OvineSNP50 BeadChip (Kijas et al., 2012). To focus on the Galway breed, a core sample set of 11 breeds, including the Galway breed, was selected for the primary population genomic analyses (*n* = 615 animals). This included populations previously examined and known to be more closely related due to their shared European origins (Howard, 2008; Kijas et al., 2012). These comparator populations also included widely used breeds, such as the Merino (MER) breed, and at-risk heritage breeds, such as the Dorset Horn (DSH), Soay (SOA) and Wiltshire (WIL) breeds (Food and Agriculture Organization, 2019) Figure 1, and Supplementary Table 1 provide further information on the geographical origins of the 11 breeds used for the core sample set analyses. In addition, Supplementary Table 1 provides information on an expanded sample set of 22 European and Asian breeds, including the core sample set, used for the phylogenetic tree and network graph reconstructions (*n* = 1,003).

**FIGURE 1.**
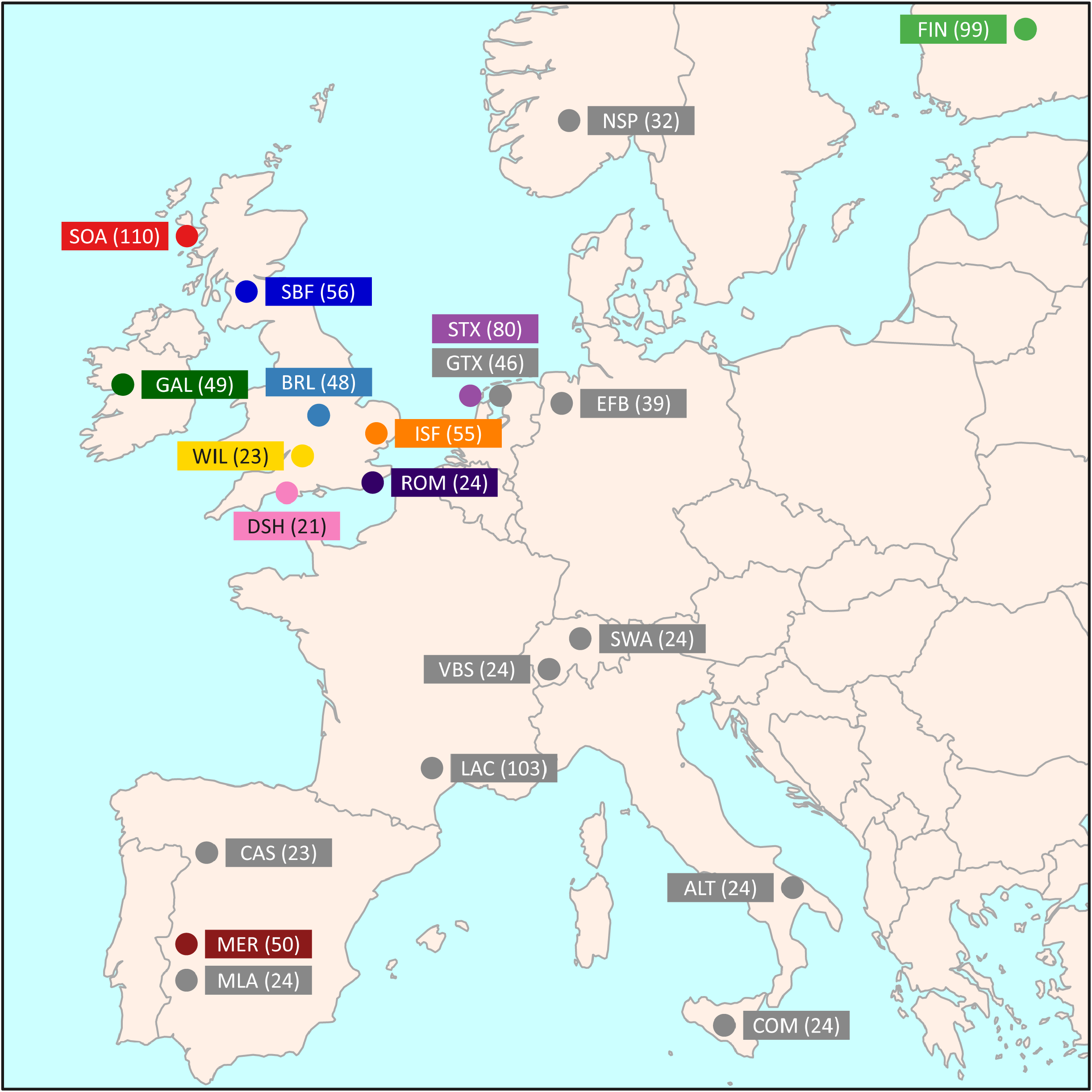
Map showing the geographical locations where breeds historically originated, adapted from Kijas et al. (2012). The number in brackets indicates the sample size. The breeds shown are the Australian Merino (MER), Border Leicester (BRL), Dorset Horn (DSH), Finnish Landrace (FIN), Galway (GAL), Irish Suffolk (ISF), New Zealand Romney (ROM), Scottish Blackface (SBF), Soay (SOA), Scottish Texel (STX) and Wiltshire (WIL).

The initial data set had already been filtered to remove SNPs with < 0.99 call rate, assay abnormality, MAF < 0.01, discordant genotypes and inheritance problems (Kijas et al., 2012). The core and extended sample genome-wide SNPs data sets for this study were filtered using PLINK v1.07 (Purcell et al., 2007) to remove SNPs lacking positional information, SNPs unassigned to any chromosome, or SNPs assigned to the X and Y chromosomes (Patterson et al., 2006; Purfield et al., 2012). The final filtered data set was composed of 47,412 SNPs with a total genotyping rate of 99.7%.

### Principal Component Analysis

Principal component analysis (PCA) was performed using 47,412 genome-wide SNPs and SMARTPCA from the EIGENSOFT software package (version 4.2) (Patterson et al., 2006). The number of autosomes was set to 26 and breed names were included. The number of outlier removal iterations was set to 0 since outliers could flag individual animals that were the result of crossbreeding. PCA plot visualisations were generated using ggplot2 (Wickham, 2016).

### *F*_ST_ Analysis

Pairwise *F*_ST_ values (Weir and Cockerham, 1984) were calculated for each pair of breeds using 47,412 genome-wide SNPs and PLINK v1.9 (Chang et al., 2015). Weighted values were chosen to account for different sample sizes for each breed.

### Construction of Phylogenetic Trees and Ancestry Graphs

Maximum likelihood (ML) phylogenetic trees with ancestry graphs were generated for the core and extended sample data sets using 47,412 genome-wide SNPs and the TreeMix (version 1.12) software package. For the core sample set, the Italian Comisana breed (COM) (Ciani et al., 2014) was used as an outgroup and five migration edges were used for TreeMix visualisation (Pickrell and Pritchard, 2012). The analysis was repeated using the extended sample set of 21 European breeds (Supplementary Table 1) and the Indian Garole breed (GAR) was used as an outgroup, again with five migration edges for TreeMix visualisation.

### Genetic Structure and Admixture History

Genetic structure and admixture history was investigated for the core sample set of the Galway and ten other breeds using 47,412 genome-wide SNPs and fastSTRUCTURE (version 1.0) (Raj et al., 2014) as described previously by us (Browett et al., 2018). The analysis was performed with the model complexity, or number of assumed populations, *K* = 2 to 11. The simple prior approach described by Raj et al. (2014) was used, which is sufficient for modelling population/breed divergence. The “true” *K*-value for the number of ancestral populations was estimated using a series of fastSTRUCTURE runs with pre-defined *K*-values that were examined using the *chooseK.py* script (Raj et al., 2014). Outputs from the fastSTRUCTURE analyses were visualised using the DISTRUCT software program (version 1.1) with standard parameters (Rosenberg, 2004).

### Modelling of Current and Historical Effective Population Size

Current and historical effective population size (*N*_e_) trends were modelled with genome-wide SNP linkage disequilibrium data from 47,412 genome-wide SNPs for the core sample set using the *SNeP* software tool (version 1.1) (Barbato et al., 2015) implementing the method for unphased SNP data as described previously by us (Browett et al., 2018). Graphs used to visualise trends in *N_e_* were generated using ggplot2 (Wickham, 2016).

### Analysis of Genomic Inbreeding and Runs of Homozygosity

Analysis of genomic inbreeding based on the inbreeding coefficient (*F*) estimated from SNP heterozygosity data was performed using 47,412 genome-wide SNPs and the PLINK v1.07 --het command (Purcell et al., 2007) since comparable inbreeding results have been observed using pruned or unpruned data for a SNP data set of similar size (Binns et al., 2012).

Runs of homozygosity (ROH) are continuous tracts of homozygosity that most likely arise due to inbreeding and can be identified through surveys of genome-wide SNP data in populations (Curik et al., 2014; Peripolli et al., 2017). Individual animal genomic inbreeding was evaluated as genome-wide autozygosity estimated from the SNP data using runs of homozygosity (ROH) values generated with PLINK v1.07 (Purcell et al., 2007) and the *F*_ROH_ statistic introduced by McQuillan et al. (2008) with methodologies previously described in detail by Purfield et al. (2012) and Browett et al. (2018). The *F*_ROH_ statistic represents the proportion of each individual animal’s genome covered by ROH, which is generally a consequence of historical inbreeding. Statistical analysis was carried out in R and graphs used to visualise *F*, *F*_ROH_ and ROH distributions were generated using ggplot2 (Wickham, 2016; R Core Team, 2018).

### Genome-wide Detection of Signatures of Selection and Functional Enrichment Analysis

The composite selection signal (CSS) method (Randhawa et al., 2014) was used to detect genomic signatures of selection as previously described (Browett et al., 2018). The CSS approach combines the fixation index (*F*_ST_), the directional change in selected allele frequency (*ΔSAF*) and cross-population extended haplotype homozygosity (XP-EHH) tests into one composite statistic for each SNP in a population genomics data set (Randhawa et al., 2014). For the present study, we used 47,412 genome-wide SNPs genotyped in 49 individual Galway sheep (GAL) samples and a sample of 50 randomly selected sheep (5 selected at random from each of the other 10 breeds in the core data set). To mitigate against false positives, genomic selection signatures were only considered significant if at least one SNP from the set of the top 0.1% genome-wide CSS scores was flanked by at least five SNPs from the set of the top 1% CSS scores.

As described previously (Browett et al., 2018), the Ensembl BioMart data mining resource (Smedley et al., 2015) was used to identify genes within ±1.0 Mb of each selection peak (Ensembl release 85, July 2016). Ingenuity^®^ Pathway Analysis (IPA^®^: Qiagen, Redwood City, CA, USA; release date July 2016) was then used to perform an overrepresentation enrichment analysis with this gene set to identify canonical pathways and functional processes of biological importance. The total gene content of Ensembl release 85 version of the OAR3.1 ovine genome assembly (Jiang et al., 2014) was used as the most appropriate reference gene set for these analyses (Timmons et al., 2015).

## RESULTS AND DISCUSSION

### Analyses of Breed Divergence, Genetic Differentiation and Admixture

The results of multiple population genomics analyses support the genetic distinctiveness of the Galway sheep population as a discrete breed. The PCA results plotted in Figure 2 demonstrate separation of the majority of breeds into distinct population clusters, with the notable exceptions of the Australian Merino (MER) and Scottish Blackface (SBF). However, it is important to note that the PCA plot visualisation shown in Figure 2 did not include the 110 samples from the Soay breed (SOA). A long history as a relatively small isolated island population (Berenos et al., 2016) has led to a marked pattern of genetic differentiation from other breeds, which is evident in the first principal component (PC1) of Supplementary Figure 1. Consequently, when the Soay breed is included in a PCA, PC3 is required to separate the Galway breed from the other populations (Supplementary Figure 2). Otherwise, the Galway breed clusters with the Scottish Texel breed (STX) and is located close to the Border Leicester breed (BLR). This result supports the documented role for the foundational New Leicester breed in the formation of the Galway and Texel breeds (Porter et al., 2016) and is compatible with the results of a previous study using autosomal microsatellites (Howard, 2008).

**FIGURE 2.**
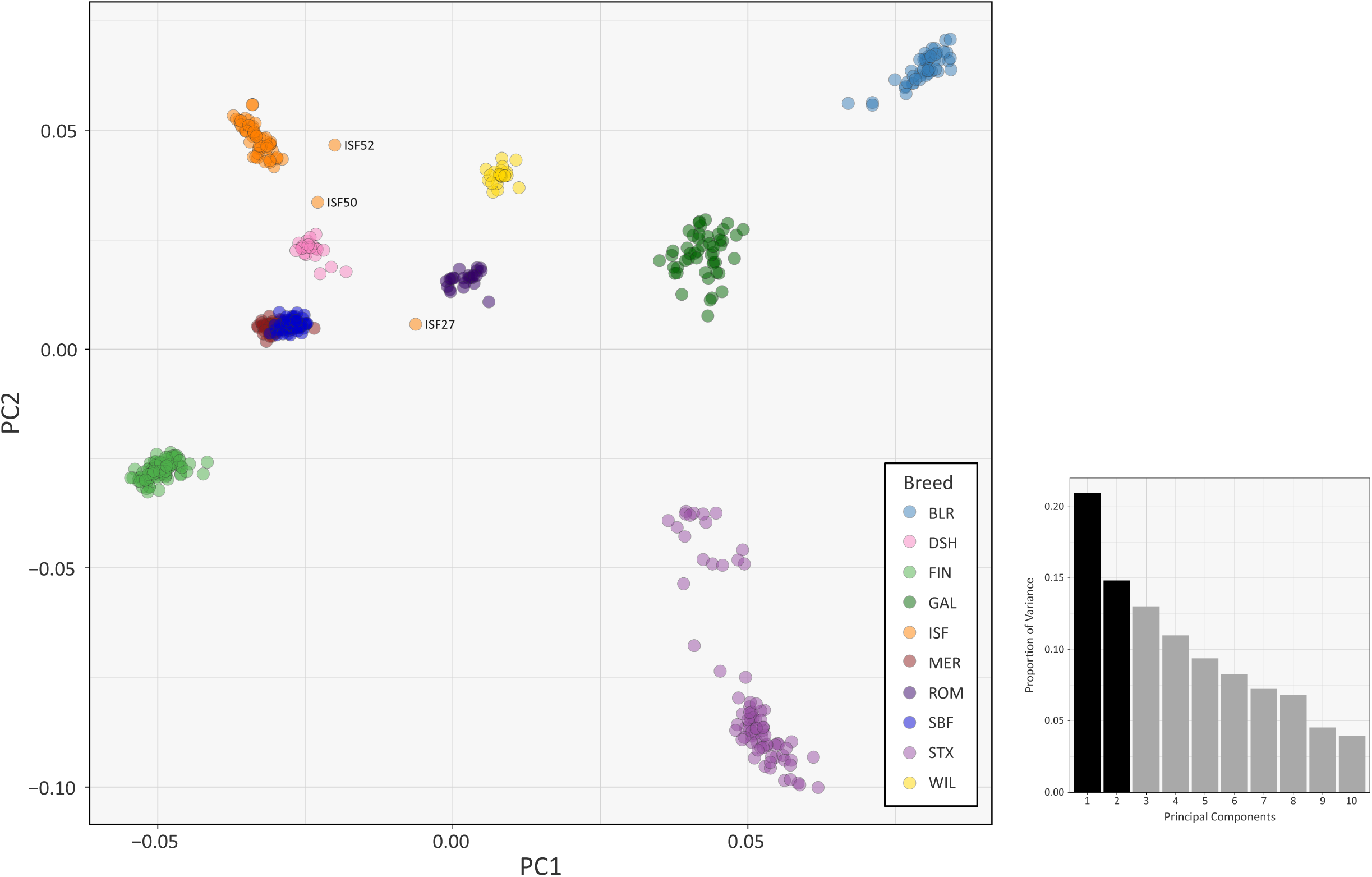
PCA plot generated using 47,412 genome-wide SNPs without the Soay sheep breed (SOA). The first principal component (PC1) is shown on the x-axis and the second principal component (PC2) is shown on the y-axis. Each breed is designated a different colour and certain individual animals that do not group by breed are labelled. The bar chart shows the proportion of variation explained by each principal component. (For comparison, PC1 versus PC3 is shown in Supplementary **Figure S3** and PC1 versus PC4 is shown in Supplementary **Figure S4**.).

The PCA plot shown in Figure 2 also demonstrated that a number of individual sheep do not cluster closely with other animals from their breeds. This is likely due to recent unacknowledged or inadvertent crossbreeding between animals from different populations (Patterson et al., 2006) or, alternatively, potential mislabelling of particular samples. For example, the 2D and 3D PCA plots shown in Supplementary Figures 1 and 2 indicate that one of the Irish Suffolk animals (ISF25) was most likely a mislabelled Scottish Texel sample as it emerged within the main Texel cluster for PC1, PC2 and PC3. Consequently, this sample ISF25 was removed from all subsequent analyses.

The PCA results are supported by the interpopulation weighted *F*_ST_ values for each pair of breeds shown in Supplementary Table 2. The results range from 0.080 (Australian Merino and Scottish Blackface) to 0.326 (Soay and Wiltshire). The pairwise *F*_ST_ values observed for the Galway population sample indicates that, with the exception of the genetically distinctive Soay sheep population (SOA), which inhabits a small island, the breed exhibits moderate genetic differentiation from other European breeds. The Galway breed exhibited relatively low pairwise *F*_ST_ values with the New Zealand Romney (ROM: 0.110), Australian Merino (MER: 0.118) and Scottish Texel (STX: 0.119) breeds. This is unsurprising because the Romney, Merino and Texel breeds are known to have shared origins with the Galway breed (Curran, 2010; Porter et al., 2016; Food and Agriculture Organization, 2019).

The ML phylogeny and ancestry graph in Figure 3 shows that the Galway breed groups closely with sheep populations of English and Dutch origin, particularly the Border Leicester (BRL) and the Scottish Texel (STX) breeds. This observation concords with previous population genomics studies (Kijas et al., 2012; Fariello et al., 2013) and known breed histories due to the shared historical input of the foundational New Leicester breed (Curran, 2010). The ML phylogeny and ancestry graph generated with additional European breeds and shown in Supplementary Figure 5 also supports the close relationship among the Galway, BRL and STX breeds. The arrows (graph edges) on Figure 3 indicate gene flow modelled between populations with the colour scale representing the weight of each migration event. Inspection of Figure 3 indicates possible historical gene flow between the Irish Suffolk and Galway branches.

**FIGURE 3.**
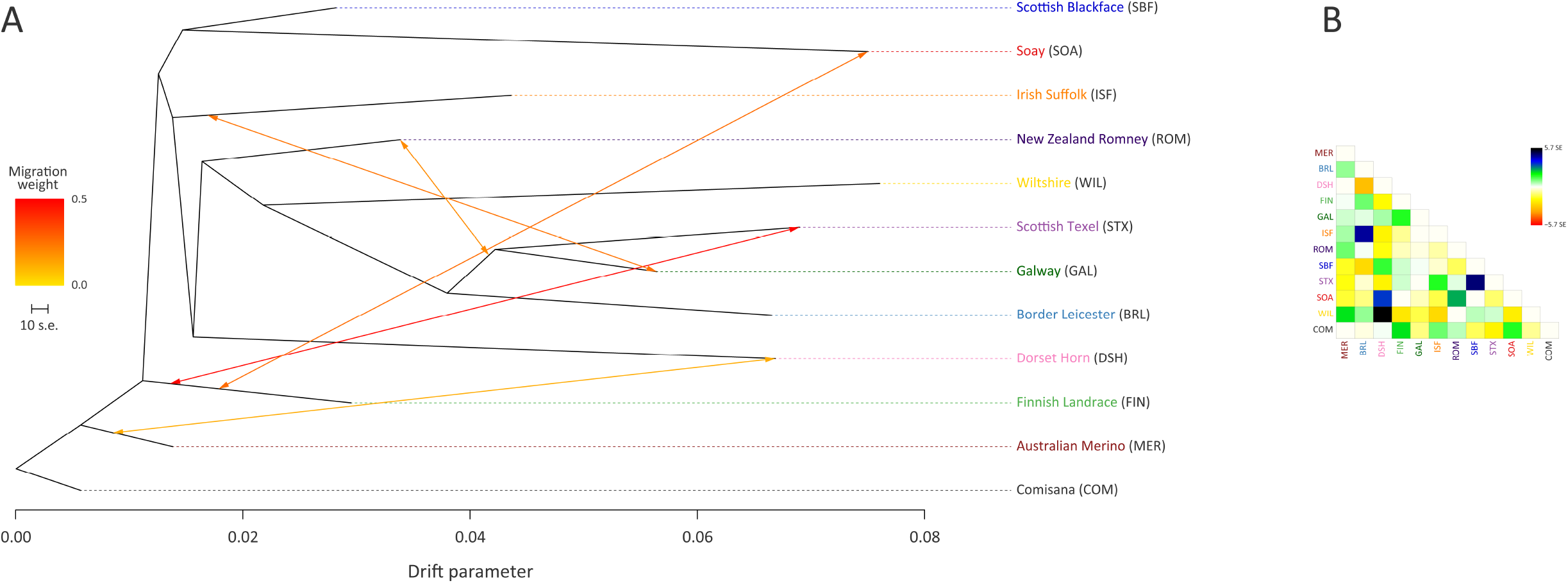
Maximum likelihood (ML) phylogenetic tree network graph generated using 47,412 genome-wide SNPs with five migration edges showing the relationships among 12 sheep breeds (A) and the residuals (B). The arrows indicate gene flow events between the populations and the colours of the arrows indicate the relative weights of migration.

Results of the genetic structure analysis for individual animals grouped by population are shown in Figure 4. Model complexity or numbers of assumed populations (*K*) ranging from 2 to 11 are visualised to explain the structure in the data and to maximise the marginal likelihood. These results demonstrate that the 11 breeds can be considered discrete populations, thereby supporting interpretation of sheep breeds as separate genetic units (Taberlet et al., 2008) and the genetic distinctiveness of Galway sheep.

**FIGURE 4.**
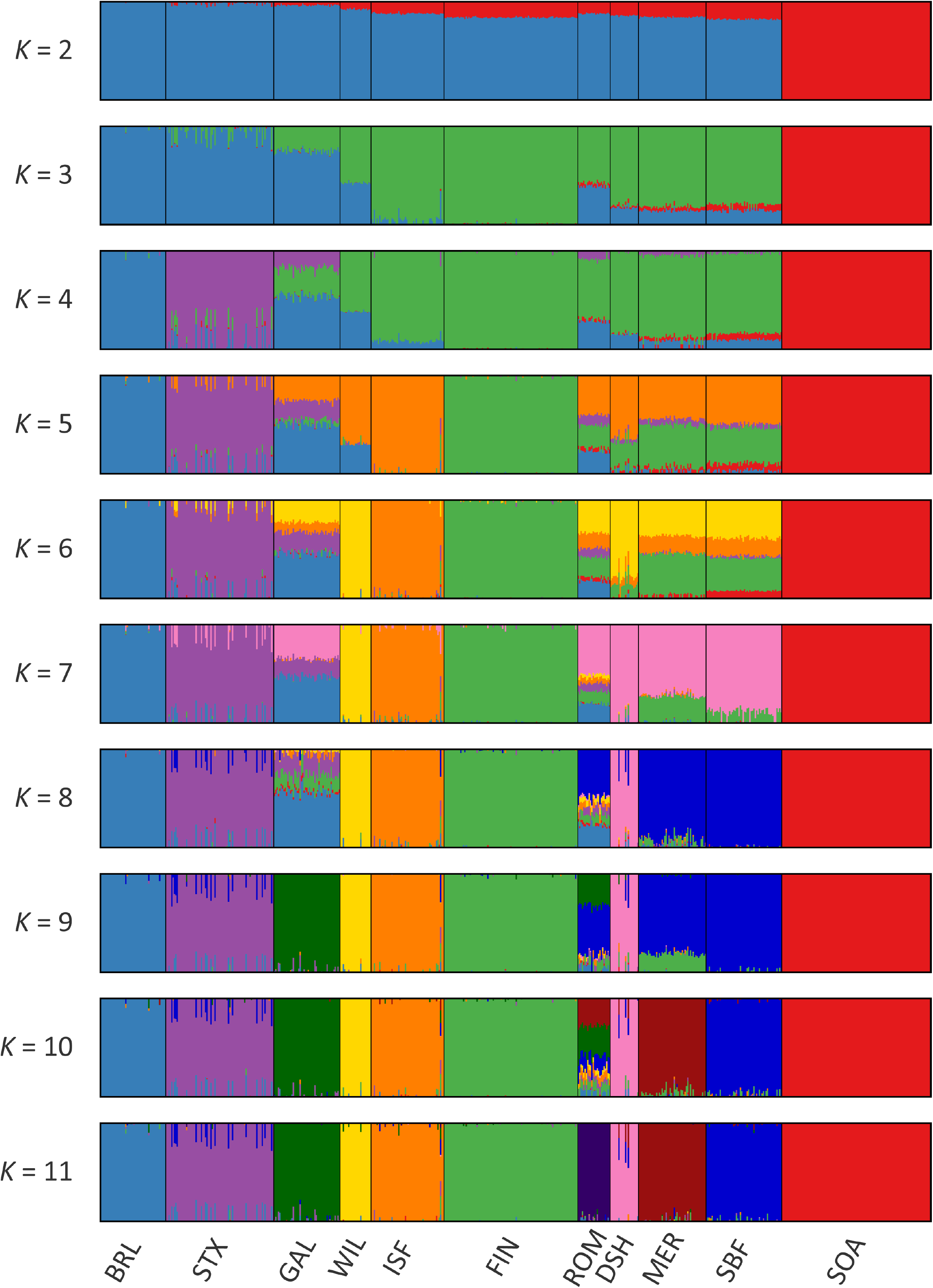
Hierarchical clustering of individual animals using 47,412 genome-wide SNPs. Results are shown for a range of assumed values (*K* = 2-11) for the number of ancestral populations.

The colours on Figure 4 indicate assignment of individual animals into modelled populations. As with the PCA shown in Supplementary Figure 1, the first split (*K* = 2) separates the isolated Soay sheep population (SOA) from the other breeds. The second split (*K* = 3) then differentiates the Finnish Landrace (FIN) from the remaining breeds. At *K* = 9 the Galway breed emerges as a distinct cluster and this genetic component is also apparent in the New Zealand Romney breed (ROM). With *K* = 11 each breed emerges as a distinct genetic cluster. However, some individual animals show evidence of prior crossbreeding or historical admixture, which is indicated by bars that exhibit varying colour proportions. Based on these results, some individual Galway animals exhibit 10% or more admixture with other sheep breeds, particularly the Border Leicester (BRL), Scottish Texel (STX) and Scottish Blackface (SBF). The observed signature of a Galway genomic component in the New Zealand Romney breed (ROM) is supported by the relatively low pairwise *F*_ST_ value for these breeds and their known origins (Supplementary Table 2) (Porter et al., 2016).

### Modelling Historical Effective Population Size

Figure 5 and Supplementary Table 4 provide the results of modelling historical effective population size (*N*_e_) for the range of conventional and at-risk sheep breeds (GAL, MER, BRL, DSH, FIN, ISF, ROM, SBF, STX and SOA). Inspection of Figure 5 and Supplementary Table 4 shows that the modelled historical trends in *N*_e_ for the 11 breeds analysed decline towards the present. However, the GAL breed are intermediate between the breeds with large census populations (FIN, ISF, MER, ROM, SBF and STX) and at-risk breeds with relatively small census populations (BRL, DSH, SOA, WIL) breeds. In addition, the most recent modelled *N*_e_ value for the GAL breed is 184 animals 13 generations ago, which is comparable to some of the breeds (e.g. ISF and STX with 178 and 150 animals, respectively). These modelled *N*_e_ values, which are based on linkage disequilibrium, may be underestimates due to the physical linkage between many SNPs (Hall, 2016).

**FIGURE 5.**
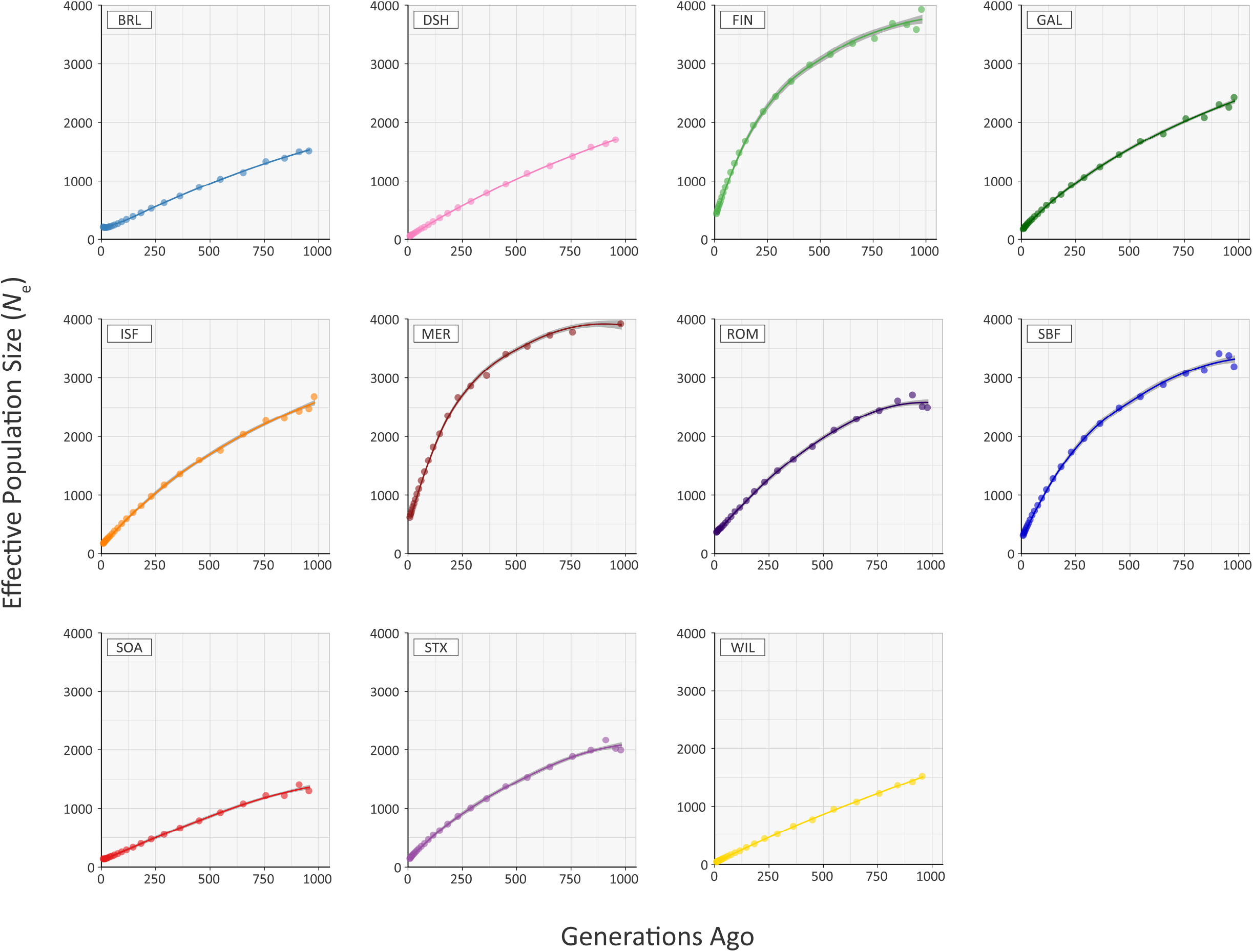
Trends in effective population size (*N*_e_) estimated using 47,412 genome-wide SNPs.

To examine these historical trends in *N*_e_ more systematically, the data for each breed were shown to be not normally distributed using the Kolmogorov-Smirnov test (Supplementary Table 3). Therefore, the non-parametric general Kruskal-Wallis test followed by pairwise Wilcoxon rank sum tests for all population/breed comparisons with adjustment for multiple statistical tests performed with the Bonferroni correction. This analysis demonstrated that the GAL historical *N*_e_ trend is significantly different only from the MER breed (*P*_adj._ = 0.006; Supplementary Table 5). Livestock populations tend to exhibit lower *N*_e_ values than comparable wild mammal populations (Waples et al., 2016). Notwithstanding this, from a conservation perspective, it is reassuring that the most recent estimated *N*_e_ value of 184 for the GAL is above the critical threshold of 100 animals considered essential for the long-term survival of livestock populations (Meuwissen, 2009). This “demographic fingerprint” (Barbato et al., 2015) is most likely a consequence of the widespread use of the Galway breed for lowland sheep production in Ireland up until the 1980s (Raftice, 2001; Curran, 2010).

### Genomic Inbreeding and Runs of Homozygosity

The recent *N*_e_ of each of the sheep breeds modelled in Figure 5 will have been substantially influenced by their inbreeding histories. In this regard, the genomic inbreeding coefficient (*F*) values estimated for individual animals across all breeds range up to 0.389 for a single Dorset Horn (DSH) animal (Figure 6). The majority of *F* values for individual animals in each breed were not normally distributed based on Shapiro-Wilk test results (Supplementary Table 3); therefore, the median *F* values were generated and evaluated for each breed (Supplementary Table 6). The breeds with the highest median *F* values were the SOA (0.308) and the WIL (0.299) and the two breeds with the lowest median *F* values were the MER (0.045) and the SBF (0.060). The other breeds exhibited intermediate median *F* values: BRL (0.243), DSH (0.169), FIN (0.087), GAL (0.127), ISF (0.185), ROM (0.086) and STX (0.111). These results provide a window on the different population histories for the breeds. For example, Soay sheep (SOA) have existed as a relatively small and isolated population on the island of Soay for hundreds of years and the Wilshere breed (WIL) have recently experienced a dramatic decline in census population and are considered at risk by the FAO (Food and Agriculture Organization, 2019). From a genetic conservation perspective, except for a single outlier (GAL26), it is encouraging that the Galway breed (GAL) exhibits an intermediate median *F* value calculated using genome-wide SNP data.

**FIGURE 6.**
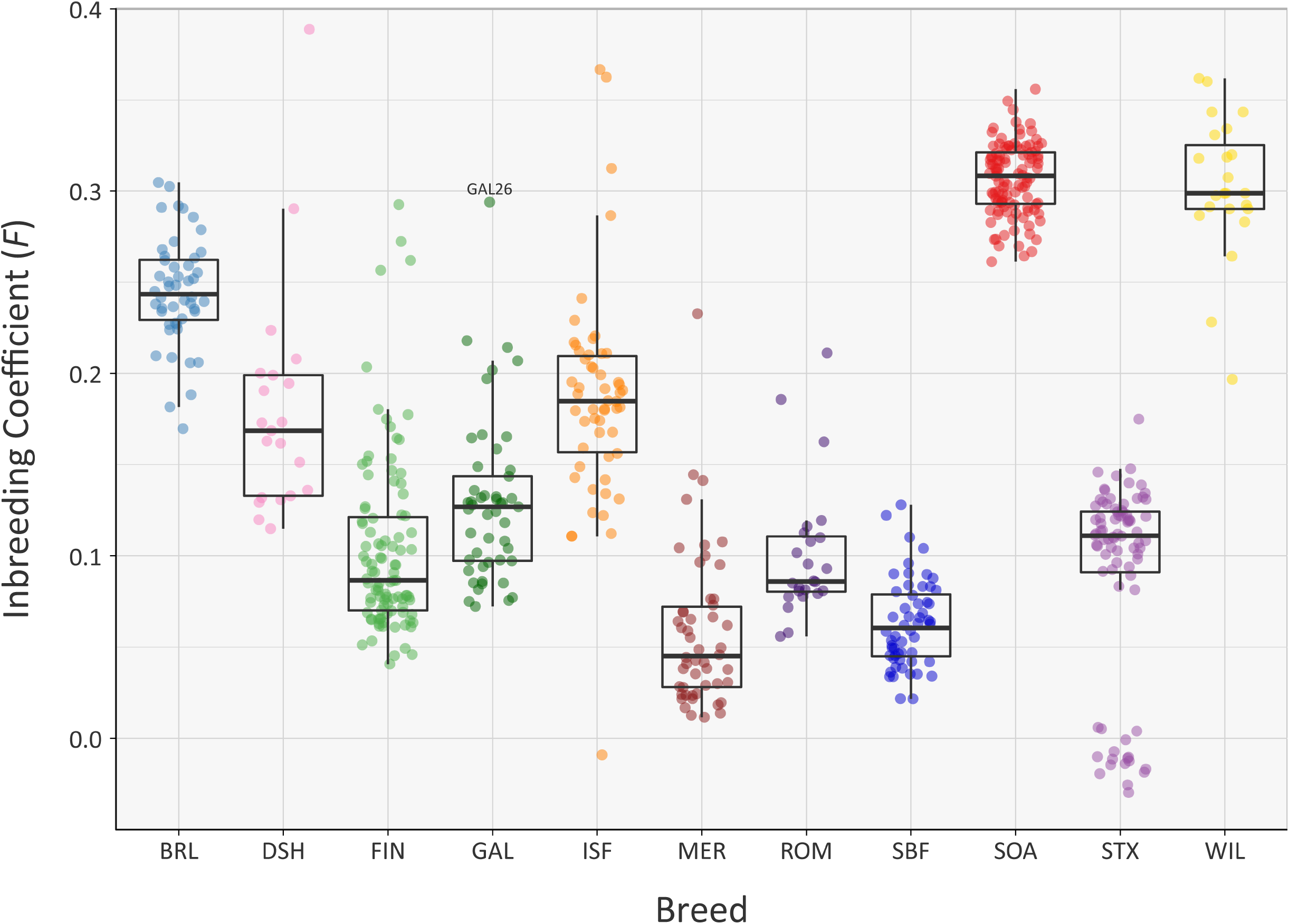
Tukey box plots showing the distribution of *F* values, estimated using 47,412 genome-wide SNPs, for the Galway sheep breed (GAL) and 10 comparator breeds. The single GAL26 outlier is labelled.

A systematic analysis of *F* value distributions using the non-parametric Kruskal-Wallis test indicated there were significant differences among breeds (*H* = 477.33, df = 10, *P* < 0.001). An analysis of all pairwise breed comparisons using the non-parametric Wilcoxon rank sum test (with Bonferroni correction) was then performed (Supplementary Table 7). These results showed that the majority of pairwise comparisons were highly significant, again reflecting the distinct demographic histories of each breed.

Overall, comparable results to those obtained using the genomic inbreeding coefficient (*F*) were observed for inbreeding coefficients estimated using ROH (*F*_ROH_) (Figure 7, Supplementary Tables 3, 6 and 8). However, there were some notable differences; in particular, the lower median *F*_ROH_ value of 0.101 for the Soay breed (SOA) is likely due to their longer geographical isolation and a consequence of early historical inbreeding that produced ROH tracts, which have broken down due to recombination (Barrett, 2012; Purfield et al., 2012). It is also notable that the Galway breed contains several individual animals with higher *F*_ROH_ values (GAL15, GAL16, GAL18, GAL26 and GAL36) indicating that this statistic is useful for identifying animals that should not be prioritised for conservation programmes. With regards to historical inbreeding in the Galway breed (GAL), inbreeding coefficients have previously been calculated using pedigree information for the population in 1969 (*F* = 0.019; Martin, 1975b), 1999 (*F* = 0.020; Raftice, 2001) and 2012 (*F* = 0.023; McHugh et al., 2014). These results indicate that the general trend in inbreeding has been relatively moderate, which may also be reflected in the results obtained using genomic information reported in the present study. It is important to note that monitoring of inbreeding for genetic conservation and management of potentially deleterious recessive genomic variants can be greatly informed through evaluation of ROH parameters using high-density SNP data (Peripolli et al., 2017)

**FIGURE 7.**
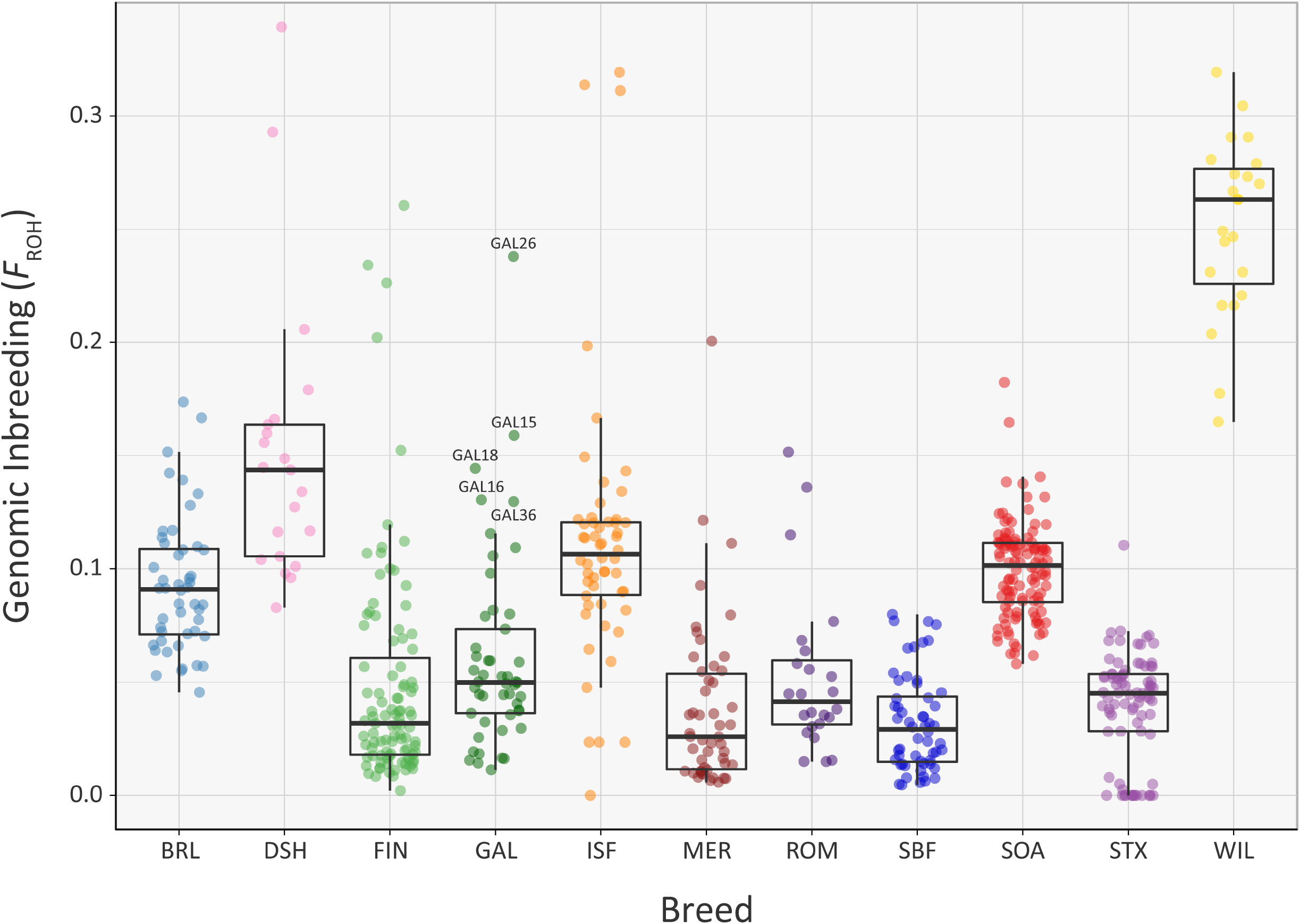
Tukey box plots showing the distribution of *F*_ROH_ values estimated using 47,412 genome-wide SNPs, for the Galway sheep breed (GAL) and 10 comparator sheep breeds. Five outlier GAL animals are labelled.

The mean sum of ROH for different length categories varies among the breeds (Figure 8); however, none of the breeds exhibit large mean values for the total length of ROH in the 1 to 5 Mb category. This is because the SNP density on the OvineSNP50 BeadChip is too low to accurately detect ROH in this size range (Purfield et al., 2012). Notwithstanding this limitation, patterns of ROH, which reflect both recent and older inbreeding histories are evident. For example, the Wiltshire breed (WIL) has large mean total ROH lengths for the other categories, presumably reflecting both historical and recent inbreeding. Other breeds, such as the Australian Merino (MER), have smaller mean total lengths of ROH in all categories, an observation that concords with the results of the genomic inbreeding and the analysis of *N*_e_ estimates. This is because individual animals from breeds with larger effective population sizes—such as the Australian Merino—are less likely to be the result of inbreeding and therefore less likely to exhibit large ROH segments in their genomes (Curik et al., 2014; Peripolli et al., 2017). The converse of this is true for breeds with lower *N_e_* values and large ROH tracts in their genomes, such as the endangered Wiltshire breed. In terms of mean total length of ROH, the Galway breed emerges between these extremes, reflecting an intermediate effective population size and history of moderate inbreeding (Figure 8). In conjunction with the other analyses of genomic diversity, these results are also encouraging for genetic conservation and the long-term viability of the breed.

**FIGURE 8.**
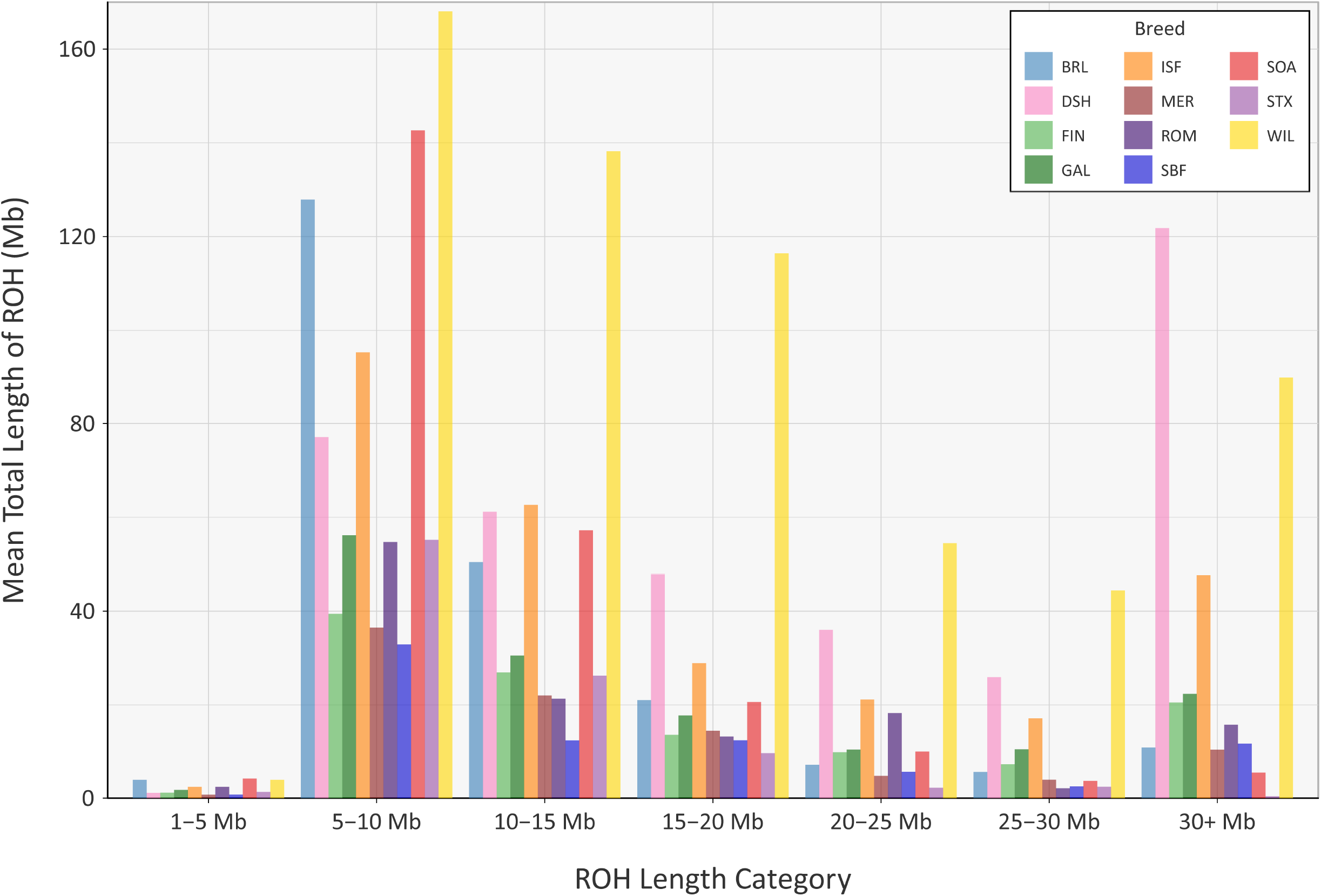
Bar graph showing the mean total length of runs of homozygosity (ROH) in different tract length categories for the Galway sheep breed (GAL) and 10 comparator sheep breeds.

### Signatures of Selection in the Galway Sheep Breed

Using defined criteria, five significant peaks of selection were detected with the CSS approach (Figure 9): two on OAR1, one on OAR3 and two on OAR8 (that merge into one peak on the graph). Each selection peak was located in a ROH tract detected in at least three Galway samples, which may indicate reduced genetic diversity in these regions as a consequence of localised selective sweeps (Purfield et al., 2017). Detection of these selection peaks demonstrates that the Galway population has experienced a unique history of both natural and human-mediated selection, presumably because of adaptation to the agroecology of Ireland, a large Northwestern European island with a temperate oceanic climate.

**FIGURE 9.**
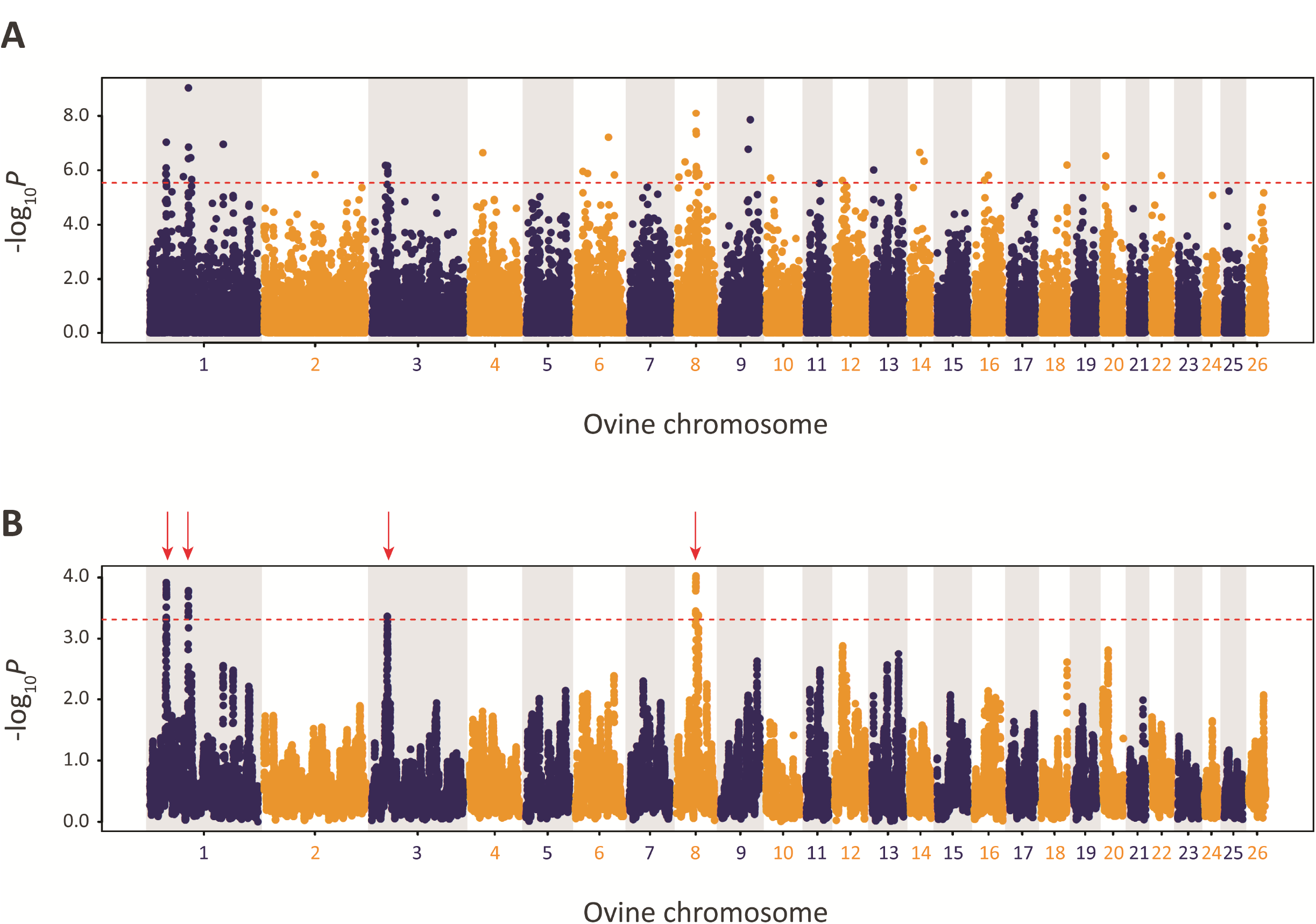
Manhattan plots of composite selection signal (CSS) results for Galway sheep (*n* = 49) contrasted with a random group selected from the other 10 breeds in the core data set (*n* = 50). (A) Unsmoothed results. (B) Smoothed results obtained by averaging CSS of SNPs within each 1Mb window. Red dotted line on each plot denotes the genome-wide 0.1% threshold for the empirical CSS scores. Red vertical arrows indicate selection peaks detected on OAR1, OAR3 and OAR8.

The precise locations of the peaks that have clusters of SNPs within the top 0.1% CSS score class are provided with additional information in Supplementary Table 9. The 197 genes within these regions are listed in Supplementary Table 10. Using IPA^®^, the top five physiological system development and function pathways enriched for the subset of 119 genes that could be mapped to HGNC symbols were identified and are listed in Table 1 (Krämer et al., 2013).

**TABLE 1.** Top five physiological system development and function pathways enriched for the 119 candidate genes proximal to the five detected selection peaks.

| Pathway | No. of Genes | Range of *P*-values |
| --- | --- | --- |
| Tissue development | 28 | 0.037–0.000 |
| Haematological system development and function | 26 | 0.037–0.000 |
| Hair and skin development and function | 7 | 0.016–0.000 |
| Immune cell trafficking | 13 | 0.037–0.001 |
| Connective tissue development and function | 15 | 0.037–0.001 |

Of the 119 candidate genes hypothesised to be under selection in the Galway breed, 28 are involved in tissue development and 15 are involved in connective tissue development and function. This is likely because the Galway breed is primarily used for meat production (Food and Agriculture Organization, 2019). In this regard, it is important to note that sheep breeds used for large-scale meat production, such as the Texel and New Zealand Romney, possess specific mutations in muscle development genes that have been subject to intense artificial selection (Cockett et al., 2005; Clop et al., 2006; Wang et al., 2016). Seven of the 119 genes are involved in hair and skin development and function, which may be explained by the use of Galway sheep in wool production (Curran, 2010). Selection and maintenance of traits that confer resilience to infectious disease is important in domestic animal populations, including many sheep breeds (Bishop and Woolliams, 2014; Bishop, 2015). Thirteen of the 119 genes under the selection peaks are involved in immune cell trafficking; which may be as a result of the climate and unique disease challenges posed by the Irish environment, such as the prevalence of liver fluke (Toolan et al., 2015). A large group of 26 genes enriched for haematological system development and function were also located under the selection peaks; however, a microevolutionary explanation for this is not hypothesised here.

### Genetic Conservation of the Galway Sheep Breed

The results of the population genomics analyses presented here are mutually consistent and highlight the utility of dense genome-wide marker data for conservation genomics in livestock populations; particularly for at-risk heritage landrace populations such as the Galway breed. Our results show the Galway breed is genetically distinct from other European sheep breeds, emerging in multivariate PCA and phylogenetic tree network graph visualisations as a distinct group but close to the Border Leicester breed (BRL), which has been observed previously (Kijas et al., 2012). In terms of effective population size and genomic inbreeding, the Galway breed emerged as intermediate between non-endangered and endangered sheep breeds. This indicates that there is substantial genetic diversity remaining in the population, which could be managed with a conservation programme that is informed by genomic information.

## DATA ACCESSIBILITY

The Galway sheep (GAL) and additional sheep breed Illumina^®^ OvineSNP50 BeadChip data are available as part of the International Sheep Genomics Consortium Ovine SNP50 HapMap Dataset (www.sheephapmap.org/download.php).

## ETHICS STATEMENT

Animal biological sample collection was conducted under license issued in accordance with Irish and European Union legislation (Cruelty to Animals Act, 1876, and European Community Directive, 86/609/EC) as described previously (Mullen et al., 2013). All animals were managed in accordance with the guidelines for the accommodation and care of animals under Article 5 of the Directive.

## AUTHOR CONTRIBUTIONS

DEM, DJH, MPM and JPH conceived and designed the project; DJH, MPM, DAM, ES and JPH organised sample collection and genotyping; GPM, SB, IAR, IWR, SDEP, MJD and CNC performed the analyses; GPM and DEM wrote the manuscript and all authors reviewed and approved the final manuscript.

## ACKNOWLEDGEMENTS

The authors would like to thank Jon Yearsley, Kay Nolan, John Leech, Lorenzo de Jonge, Charlotte Gilbert and Han Haige for helpful discussion and feedback. We also thank members of the Galway Sheep Breeders Association for their cooperation in sample and data collection. Genotypic data used in this study was collected as part of a coordinated international project run under the auspices of the International Sheep Genomics Consortium.

## REFERENCES

1. Allendorf, FW, Luikart, G, and Aitken, SN (2013). Conservation and the Genetics of Populations. Oxford, UK: John Wiley & Sons.

2. Barbato, M, Orozco-terWengel, P, Tapio, M, and Bruford, MW (2015). *SNeP*: a tool to estimate trends in recent effective population size trajectories using genome-wide SNP data. Front Genet 6, 109. doi: 10.3389/fgene.2015.00109.

3. Barrett, RD (2012). Bad coat, ripped genes: cryptic selection on coat colour varies with ontogeny in Soay sheep. Mol Ecol 21, 2833–2835. doi: 10.1111/j.1365-294X.2012.05560.x.

4. Berenos, C, Ellis, PA, Pilkington, JG, and Pemberton, JM (2016). Genomic analysis reveals depression due to both individual and maternal inbreeding in a free-living mammal population. Mol Ecol 25, 3152–3168. doi: 10.1111/mec.13681.

5. Binns, MM, Boehler, DA, Bailey, E, Lear, TL, Cardwell, JM, and Lambert, DH (2012). Inbreeding in the Thoroughbred horse. Anim Genet 43, 340–342. doi: 10.1111/j.1365-2052.2011.02259.x.

6. Biscarini, F, Nicolazzi, EL, Stella, A, Boettcher, PJ, and Gandini, G (2015). Challenges and opportunities in genetic improvement of local livestock breeds. Front Genet 6, 33. doi: 10.3389/fgene.2015.00033.

7. Bishop, SC (2015). Genetic resistance to infections in sheep. Vet Microbiol 181, 2–7. doi: 10.1016/j.vetmic.2015.07.013.

8. Bishop, SC, and Woolliams, JA (2014). Genomics and disease resistance studies in livestock. Livestock Sci 166, 190–198. doi: 10.1016/j.livsci.2014.04.034.

9. Bowles, D (2015). Recent advances in understanding the genetic resources of sheep breeds locally-adapted to the UK uplands: opportunities they offer for sustainable productivity. Front Genet 6, 24. doi: 10.3389/fgene.2015.00024.

10. Browett, S, McHugo, G, Richardson, IW, Magee, DA, Park, SDE, Fahey, AG, Kearney, JF, Correia, CN, Randhawa, IAS, and MacHugh, DE (2018). Genomic characterisation of the indigenous Irish Kerry cattle breed. Front Genet 9, 51. doi: 10.3389/fgene.2018.00051.

11. Chang, CC, Chow, CC, Tellier, LC, Vattikuti, S, Purcell, SM, and Lee, JJ (2015). Second-generation PLINK: rising to the challenge of larger and richer datasets. Gigascience 4, 7. doi: 10.1186/s13742-015-0047-8.

12. Ciani, E, Crepaldi, P, Nicoloso, L, Lasagna, E, Sarti, FM, Moioli, B, Napolitano, F, Carta, A, Usai, G, D’Andrea, M, Marletta, D, Ciampolini, R, Riggio, V, Occidente, M, Matassino, D, Kompan, D, Modesto, P, Macciotta, N, Ajmone-Marsan, P, and Pilla, F (2014). Genome-wide analysis of Italian sheep diversity reveals a strong geographic pattern and cryptic relationships between breeds. Anim Genet 45, 256–266. doi: 10.1111/age.12106.

13. Clop, A, Marcq, F, Takeda, H, Pirottin, D, Tordoir, X, Bibe, B, Bouix, J, Caiment, F, Elsen, JM, Eychenne, F, Larzul, C, Laville, E, Meish, F, Milenkovic, D, Tobin, J, Charlier, C, and Georges, M (2006). A mutation creating a potential illegitimate microRNA target site in the myostatin gene affects muscularity in sheep. Nat Genet 38, 813–818.

14. Cockett, NE, Smit, MA, Bidwell, CA, Segers, K, Hadfield, TL, Snowder, GD, Georges, M, and Charlier, C (2005). The callipyge mutation and other genes that affect muscle hypertrophy in sheep. Genet Sel Evol 37 Suppl 1, S65–81.

15. Curik, I, Ferenčaković, M, and Sölkner, J (2014). Inbreeding and runs of homozygosity: A possible solution to an old problem. Livestock Sci 166, 26–34. doi: 10.1016/j.livsci.2014.05.034.

16. Curran, PL (2010). The Native Lowland Sheep of Galway and Roscommon: a History. Tara, Co. Meath, Ireland: Patrick Leonard Curran.

17. Fariello, MI, Boitard, S, Naya, H, SanCristobal, M, and Servin, B (2013). Detecting signatures of selection through haplotype differentiation among hierarchically structured populations. Genetics 193, 929–941. doi: 10.1534/genetics.112.147231.

18. Food and Agriculture Organization. 2019. Domestic Animal Diversity Information System (DAD-IS); available at: www.fao.org/dadis. [Accessed 14/02/2019].

19. Hall, SJ (2016). Effective population sizes in cattle, sheep, horses, pigs and goats estimated from census and herdbook data. Animal 10, 1778–1785. doi: 10.1017/s1751731116000914.

20. Hanrahan, JP. 1999. The Galway Breed ‒ Origins and the Future; available at www.galwaysheep.ie. [Accessed 13/07/2018].

21. Howard, DJ (2008). A Comparative Study of Molecular Genetic Variation in Three Irish Sheep Breeds; the Galway, Texel and Suffolk. PhD thesis, University College Dublin.

22. Jiang, Y, Xie, M, Chen, W, Talbot, R, Maddox, JF, Faraut, T, Wu, C, Muzny, DM, Li, Y, Zhang, W, Stanton, JA, Brauning, R, Barris, WC, Hourlier, T, Aken, BL, Searle, SM, Adelson, DL, Bian, C, Cam, GR, Chen, Y, Cheng, S, DeSilva, U, Dixen, K, Dong, Y, Fan, G, Franklin, IR, Fu, S, Fuentes-Utrilla, P, Guan, R, Highland, MA, Holder, ME, Huang, G, Ingham, AB, Jhangiani, SN, Kalra, D, Kovar, CL, Lee, SL, Liu, W, Liu, X, Lu, C, Lv, T, Mathew, T, McWilliam, S, Menzies, M, Pan, S, Robelin, D, Servin, B, Townley, D, Wang, W, Wei, B, White, SN, Yang, X, Ye, C, Yue, Y, Zeng, P, Zhou, Q, Hansen, JB, Kristiansen, K, Gibbs, RA, Flicek, P, Warkup, CC, Jones, HE, Oddy, VH, Nicholas, FW, McEwan, JC, Kijas, JW, Wang, J, Worley, KC, Archibald, AL, Cockett, N, Xu, X, Wang, W, and Dalrymple, BP (2014). The sheep genome illuminates biology of the rumen and lipid metabolism. Science 344, 1168–1173. doi: 10.1126/science.1252806.

23. Kijas, JW, Lenstra, JA, Hayes, B, Boitard, S, Porto Neto, LR, San Cristobal, M, Servin, B, McCulloch, R, Whan, V, Gietzen, K, Paiva, S, Barendse, W, Ciani, E, Raadsma, H, McEwan, J, Dalrymple, B, and International Sheep Genomics Consortium (2012). Genome-wide analysis of the world’s sheep breeds reveals high levels of historic mixture and strong recent selection. PLoS Biol 10, e1001258. doi: 10.1371/journal.pbio.1001258.

24. Kijas, JW, Townley, D, Dalrymple, BP, Heaton, MP, Maddox, JF, McGrath, A, Wilson, P, Ingersoll, RG, McCulloch, R, McWilliam, S, Tang, D, McEwan, J, Cockett, N, Oddy, VH, Nicholas, FW, and Raadsma, H (2009). A genome wide survey of SNP variation reveals the genetic structure of sheep breeds. PLoS ONE 4, e4668. doi: 10.1371/journal.pone.0004668.

25. Krämer, A, Green, J, Pollard, J Jr, and Tugendreich, S (2013). Causal analysis approaches in ingenuity pathway analysis. Bioinformatics 30, 523–530.

26. Larson, G, and Fuller, DQ (2014). The evolution of animal domestication. Annu Rev Ecol Evol Syst 45, 115–136. doi: 10.1146/annurev-ecolsys-110512-135813.

27. MacHugh, DE, Larson, G, and Orlando, L (2017). Taming the past: ancient DNA and the study of animal domestication. Annu Rev Anim Biosci 5, 329–351. doi: 10.1146/annurev-animal-022516-022747.

28. Martin, I (1975a). A genetic analysis of the Galway sheep breed: 1. some aspects of population dynamics of the pedigree and non-pedigree Galway sheep breed. Ir J Agric Sci 14, 245–253.

29. Martin, I (1975b). A genetic analysis of the Galway sheep breed: 3. level of inbreeding in the pedigree Galway sheep breed. Ir J Agric Sci 14, 269–274.

30. McHugh, N, Berry, D, McParland, S, Wall, E, and Pabiou, T (2014). Irish sheep breeding: current status and future plans. Teagasc and Sheep Ireland.

31. McQuillan, R, Leutenegger, AL, Abdel-Rahman, R, Franklin, CS, Pericic, M, Barac-Lauc, L, Smolej-Narancic, N, Janicijevic, B, Polasek, O, Tenesa, A, Macleod, AK, Farrington, SM, Rudan, P, Hayward, C, Vitart, V, Rudan, I, Wild, SH, Dunlop, MG, Wright, AF, Campbell, H, and Wilson, JF (2008). Runs of homozygosity in European populations. Am J Hum Genet 83, 359–372. doi: 10.1016/j.ajhg.2008.08.007.

32. Meuwissen, T (2009). Genetic management of small populations: A review. Acta Agric Scand A Anim Sci 59, 71–79. doi: 10.1080/09064700903118148.

33. Mullen, MP, Hanrahan, JP, Howard, DJ, and Powell, R (2013). Investigation of prolific sheep from UK and Ireland for evidence on origin of the mutations in *BMP15* (*Fec*X*^G^*, *Fec*X*^B^*) and *GDF9* (*Fec*G*^H^*) in Belclare and Cambridge sheep. PLoS ONE 8, e53172. doi: 10.1371/journal.pone.0053172.

34. Patterson, N, Price, AL, and Reich, D (2006). Population structure and eigenanalysis. PLoS Genet 2, e190. doi: 10.1371/journal.pgen.0020190.

35. Peripolli, E, Munari, DP, Silva, M, Lima, ALF, Irgang, R, and Baldi, F (2017). Runs of homozygosity: current knowledge and applications in livestock. Anim Genet 48, 255–271. doi: 10.1111/age.12526.

36. Petersen, B (2017). Basics of genome editing technology and its application in livestock species. Reprod Domest Anim 52 Suppl 3, 4–13. doi: 10.1111/rda.13012.

37. Pickrell, JK, and Pritchard, JK (2012). Inference of population splits and mixtures from genome-wide allele frequency data. PLoS Genet 8, e1002967. doi: 10.1371/journal.pgen.1002967.

38. Porter, V, Alderson, L, Hall, SJG, and Sponenberg, DP (2016). Mason’s World Encyclopedia of Livestock Breeds and Breeding. Wallingford, UK: CABI.

39. Purcell, S, Neale, B, Todd-Brown, K, Thomas, L, Ferreira, MA, Bender, D, Maller, J, Sklar, P, de Bakker, PI, Daly, MJ, and Sham, PC (2007). PLINK: a tool set for whole-genome association and population-based linkage analyses. Am J Hum Genet 81, 559–575. doi: 10.1086/519795.

40. Purfield, DC, Berry, DP, McParland, S, and Bradley, DG (2012). Runs of homozygosity and population history in cattle. BMC Genet 13, 70. doi: 10.1186/1471-2156-13-70.

41. Purfield, DC, McParland, S, Wall, E, and Berry, DP (2017). The distribution of runs of homozygosity and selection signatures in six commercial meat sheep breeds. PLoS ONE 12, e0176780. doi: 10.1371/journal.pone.0176780.

42. R Core Team. 2018. R: A Language and Environment for Statistical Computing. Available: https://www.R-project.org.

43. Raftice, MJ (2001). Genetic Conservation of the Galway Sheep Breed. MSc thesis, University College Dublin.

44. Raj, A, Stephens, M, and Pritchard, JK (2014). fastSTRUCTURE: variational inference of population structure in large SNP data sets. Genetics 197, 573–589. doi: 10.1534/genetics.114.164350.

45. Randhawa, IA, Khatkar, MS, Thomson, PC, and Raadsma, HW (2014). Composite selection signals can localize the trait specific genomic regions in multi-breed populations of cattle and sheep. BMC Genet 15, 34. doi: 10.1186/1471-2156-15-34.

46. Rosenberg, NA (2004). DISTRUCT: a program for the graphical display of population structure. Mol Ecol Notes 4, 137–138.

47. Smedley, D, Haider, S, Durinck, S, Pandini, L, Provero, P, Allen, J, Arnaiz, O, Awedh, MH, Baldock, R, Barbiera, G, Bardou, P, Beck, T, Blake, A, Bonierbale, M, Brookes, AJ, Bucci, G, Buetti, I, Burge, S, Cabau, C, Carlson, JW, Chelala, C, Chrysostomou, C, Cittaro, D, Collin, O, Cordova, R, Cutts, RJ, Dassi, E, Di Genova, A, Djari, A, Esposito, A, Estrella, H, Eyras, E, Fernandez-Banet, J, Forbes, S, Free, RC, Fujisawa, T, Gadaleta, E, Garcia-Manteiga, JM, Goodstein, D, Gray, K, Guerra-Assuncao, JA, Haggarty, B, Han, DJ, Han, BW, Harris, T, Harshbarger, J, Hastings, RK, Hayes, RD, Hoede, C, Hu, S, Hu, ZL, Hutchins, L, Kan, Z, Kawaji, H, Keliet, A, Kerhornou, A, Kim, S, Kinsella, R, Klopp, C, Kong, L, Lawson, D, Lazarevic, D, Lee, JH, Letellier, T, Li, CY, Lio, P, Liu, CJ, Luo, J, Maass, A, Mariette, J, Maurel, T, Merella, S, Mohamed, AM, Moreews, F, Nabihoudine, I, Ndegwa, N, Noirot, C, Perez-Llamas, C, Primig, M, Quattrone, A, Quesneville, H, Rambaldi, D, Reecy, J, Riba, M, Rosanoff, S, Saddiq, AA, Salas, E, Sallou, O, Shepherd, R, Simon, R, Sperling, L, Spooner, W, Staines, DM, Steinbach, D, Stone, K, Stupka, E, Teague, JW, Dayem Ullah, AZ, Wang, J, Ware, D, et al. (2015). The BioMart community portal: an innovative alternative to large, centralized data repositories. Nucleic Acids Res 43, W589–598. doi: 10.1093/nar/gkv350.

48. Taberlet, P, Coissac, E, Pansu, J, and Pompanon, F (2011). Conservation genetics of cattle, sheep, and goats. C R Biol 334, 247–254. doi: 10.1016/j.crvi.2010.12.007.

49. Taberlet, P, Valentini, A, Rezaei, H, Naderi, S, Pompanon, F, Negrini, R, and Ajmone-Marsan, P (2008). Are cattle, sheep, and goats endangered species? Mol Ecol 17, 275–284.

50. Tapio, M, Tapio, I, Grislis, Z, Holm, LE, Jeppsson, S, Kantanen, J, Miceikiene, I, Olsaker, I, Viinalass, H, and Eythorsdottir, E (2005). Native breeds demonstrate high contributions to the molecular variation in northern European sheep. Mol Ecol 14, 3951–3963. doi: 10.1111/j.1365-294X.2005.02727.x.

51. Timmons, JA, Szkop, KJ, and Gallagher, IJ (2015). Multiple sources of bias confound functional enrichment analysis of global-omics data. Genome Biol 16, 186. doi: 10.1186/s13059-015-0761-7.

52. Toolan, DP, Mitchell, G, Searle, K, Sheehan, M, Skuce, PJ, and Zadoks, RN (2015). Bovine and ovine rumen fluke in Ireland—Prevalence, risk factors and species identity based on passive veterinary surveillance and abattoir findings. Vet Parasitol 212, 168–174.

53. Van Eenennaam, AL (2017). Genetic modification of food animals. Curr Opin Biotechnol 44, 27–34. doi: http://dx.doi.org/10.1016/j.copbio.2016.10.007.

54. Wang, J, Zhou, H, Hu, J, Li, S, Luo, Y, and Hickford, JG (2016). Two single nucleotide polymorphisms in the promoter of the ovine myostatin gene (*MSTN*) and their effect on growth and carcass muscle traits in New Zealand Romney sheep. J Anim Breed Genet 133, 219–226. doi: 10.1111/jbg.12171.

55. Waples, RK, Larson, WA, and Waples, RS (2016). Estimating contemporary effective population size in non-model species using linkage disequilibrium across thousands of loci. Heredity (Edinb) 117, 233–240. doi: 10.1038/hdy.2016.60.

56. Weir, BS, and Cockerham, CC (1984). Estimating F-statistics for the analysis of population structure. Evolution, 1358–1370.

57. Wells, KD (2013). Natural genotypes via genetic engineering. Proc Natl Acad Sci U S A 110, 16295–16296. doi: 10.1073/pnas.1315623110.

58. Wickham, H (2016). ggplot2: Elegant Graphics for Data Analysis. Springer Publishing Company, New York, USA.

59. Wykes, DL (2004). Robert Bakewell (1725-1795) of Dishley: farmer and livestock improver. Agric Hist Rev 52, 38–55.

