## Supplementary Material for "A Population Genomics Analysis of the Native Irish Galway Sheep Breed"

### 1 Supplementary Figures and Tables

#### 1.1. Supplementary Figures

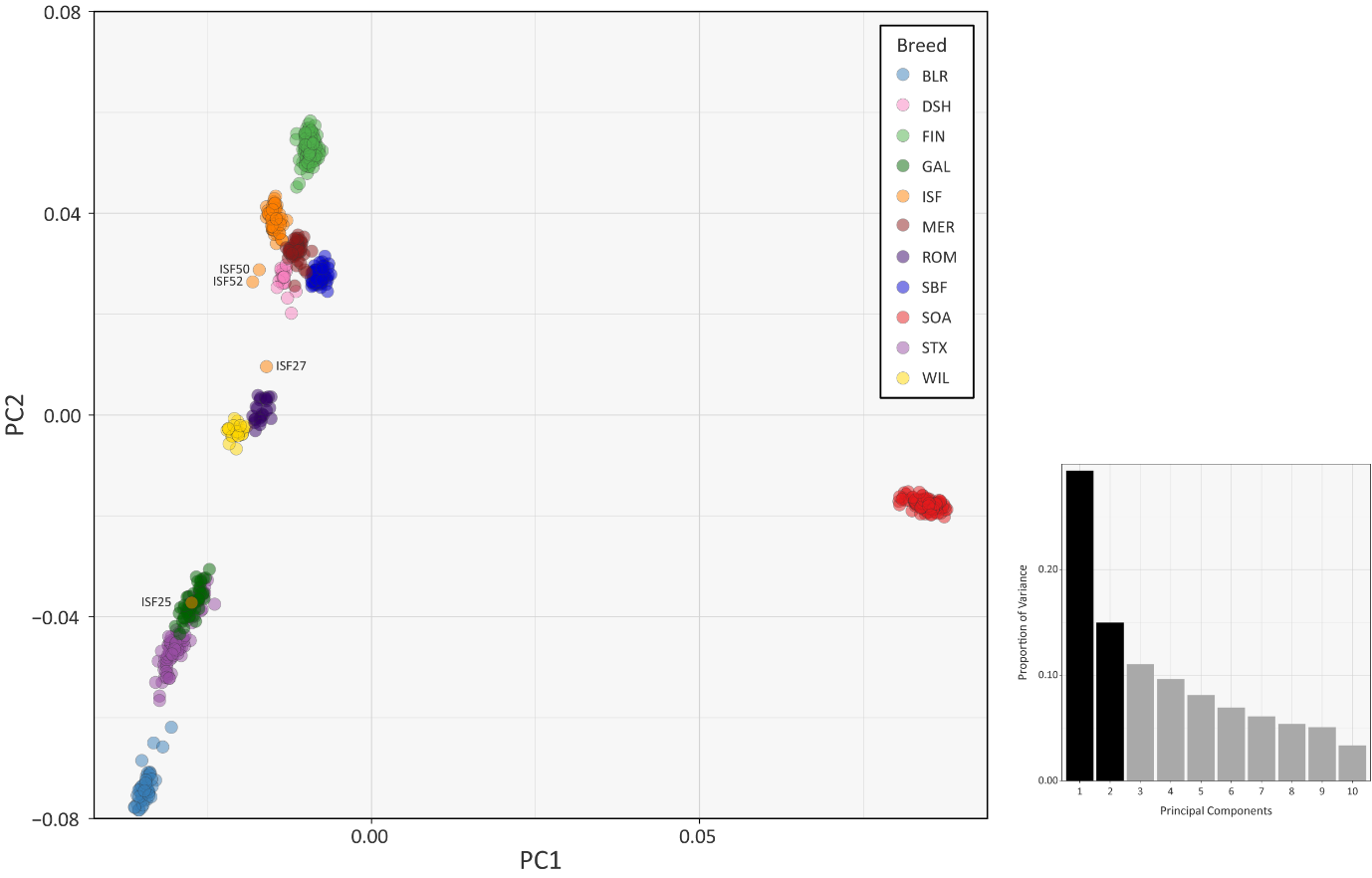

**Supplementary Figure 1.** PCA plot generated using 47,412 genome-wide SNPs. Each breed is designated by a different colour and individuals that do not group with the rest of their breed are labelled. The bar chart shows the percentage of variation explained by each principal component. ISF25 is most likely a mislabelled sample.

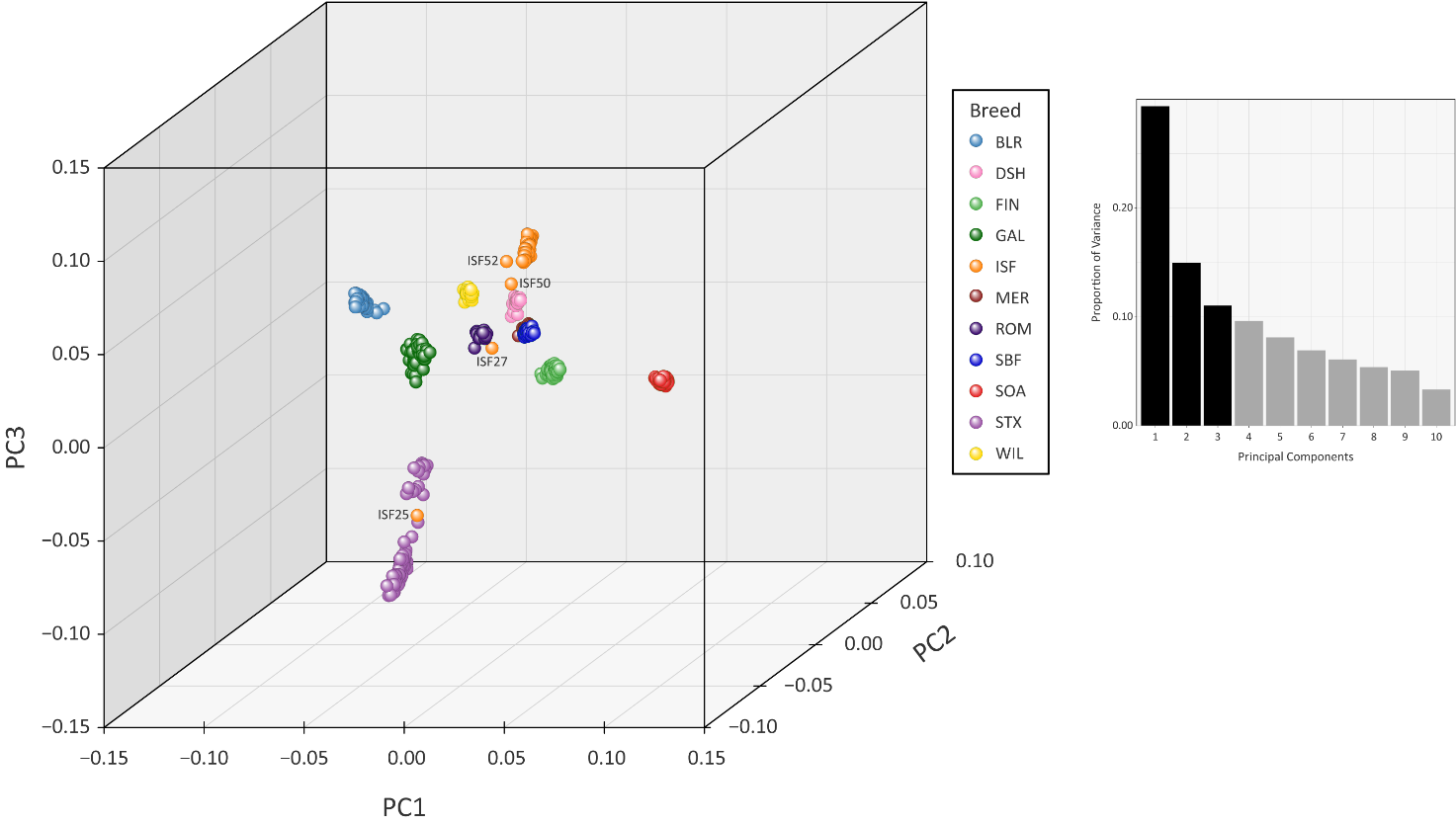

**Supplementary Figure 2.** PCA plot generated using 47,412 genome-wide SNPs. Each breed is designated by a different colour and individual animals that do not group by breed are labelled. The bar chart shows the proportion of variation explained by the first 10 principal components. ISF25 is most likely a mislabelled sample.

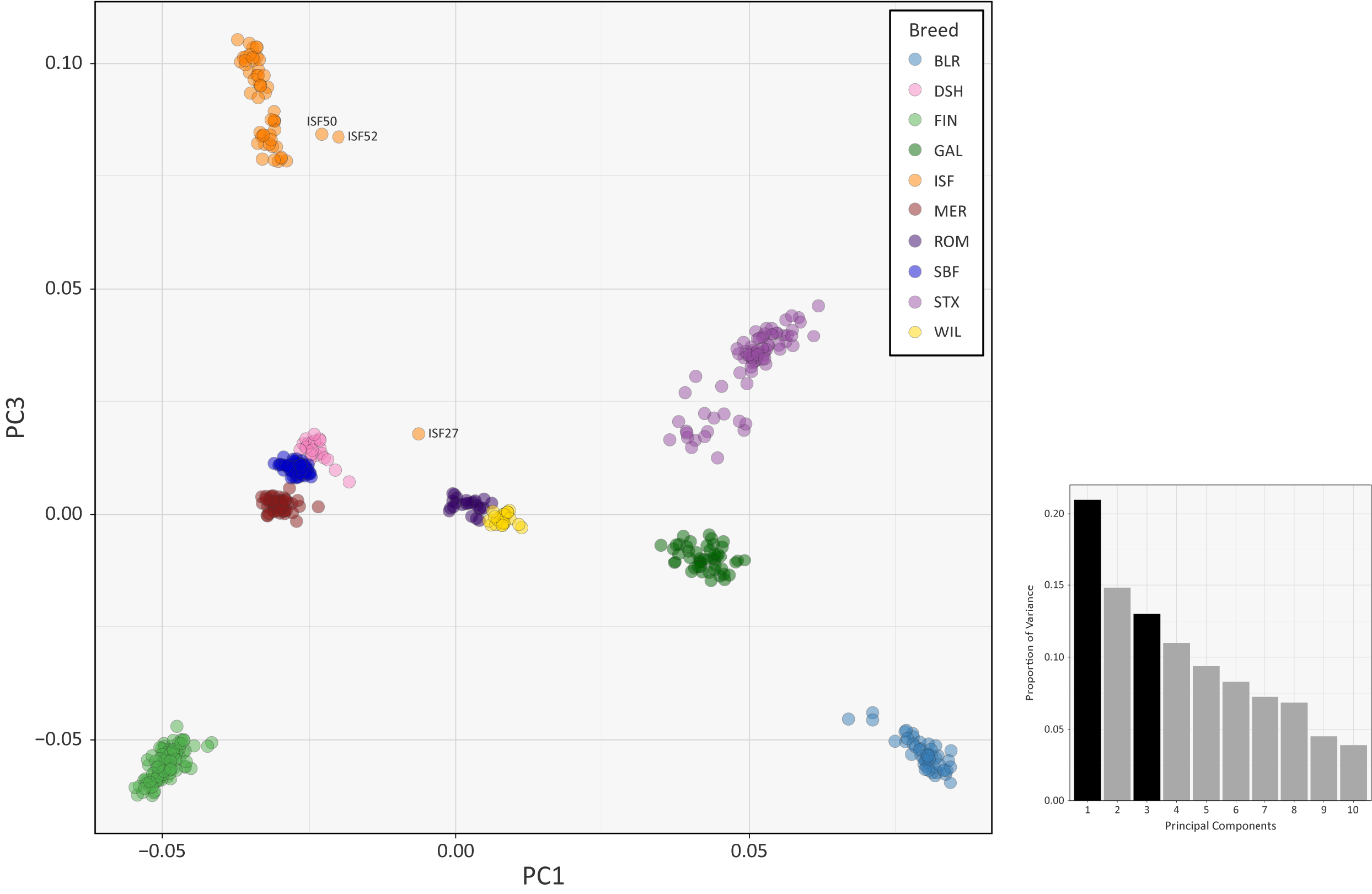

**Supplementary Figure 3.** PCA plot generated using 47,412 genome-wide SNPs without the Soay sheep breed (SOA). Each breed is designated by a different colour and individual animals that do not group by breed are labelled. The bar chart shows the proportion of variation explained by the first 10 principal components.

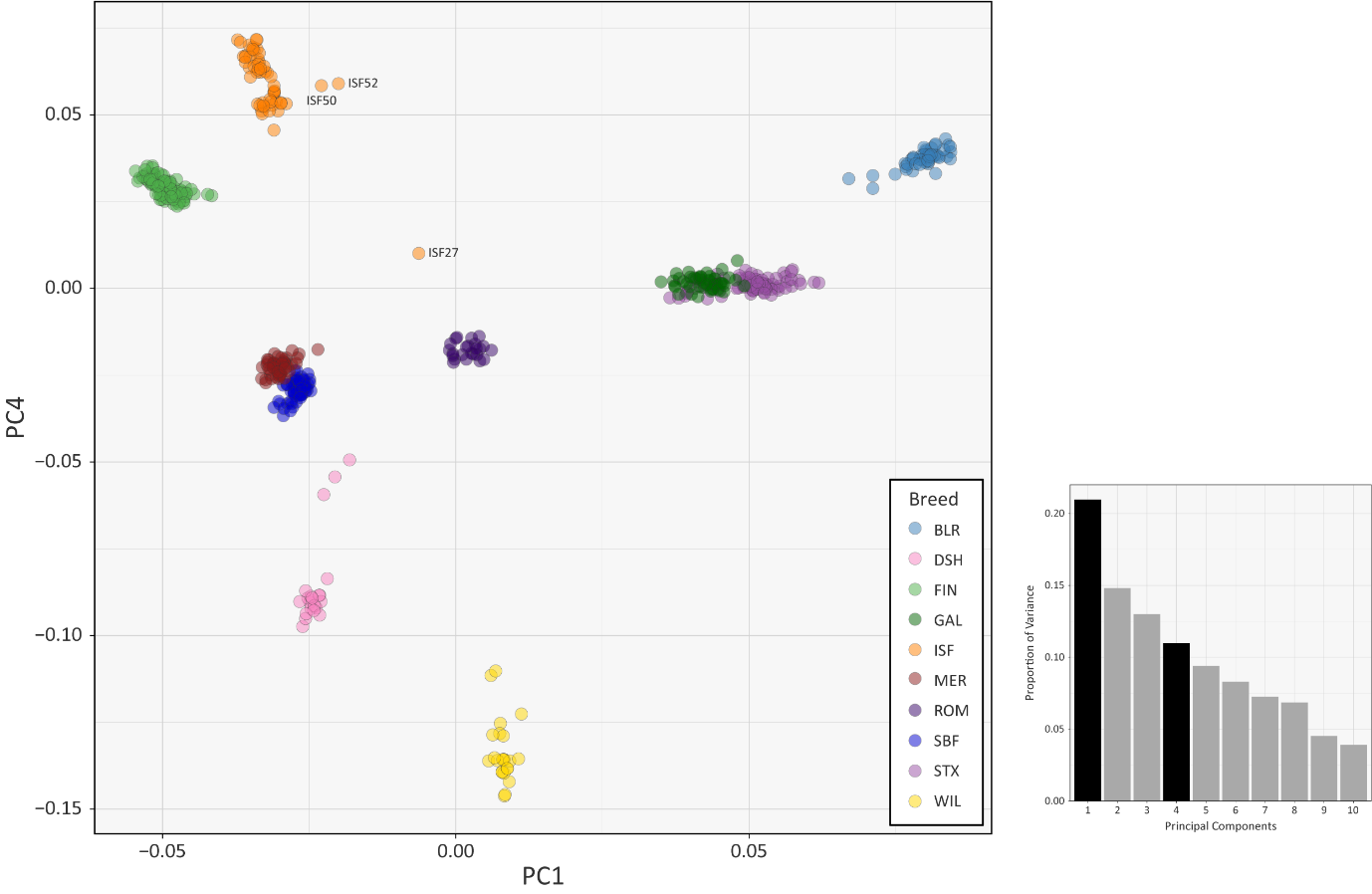

**Supplementary Figure 4.** PCA plot generated using 47,412 genome-wide SNPs without the Soay sheep breed (SOA). Each breed is designated by a different colour and individual animals that do not group by breed are labelled. The bar chart shows the proportion of variation explained by the first 10 principal components.

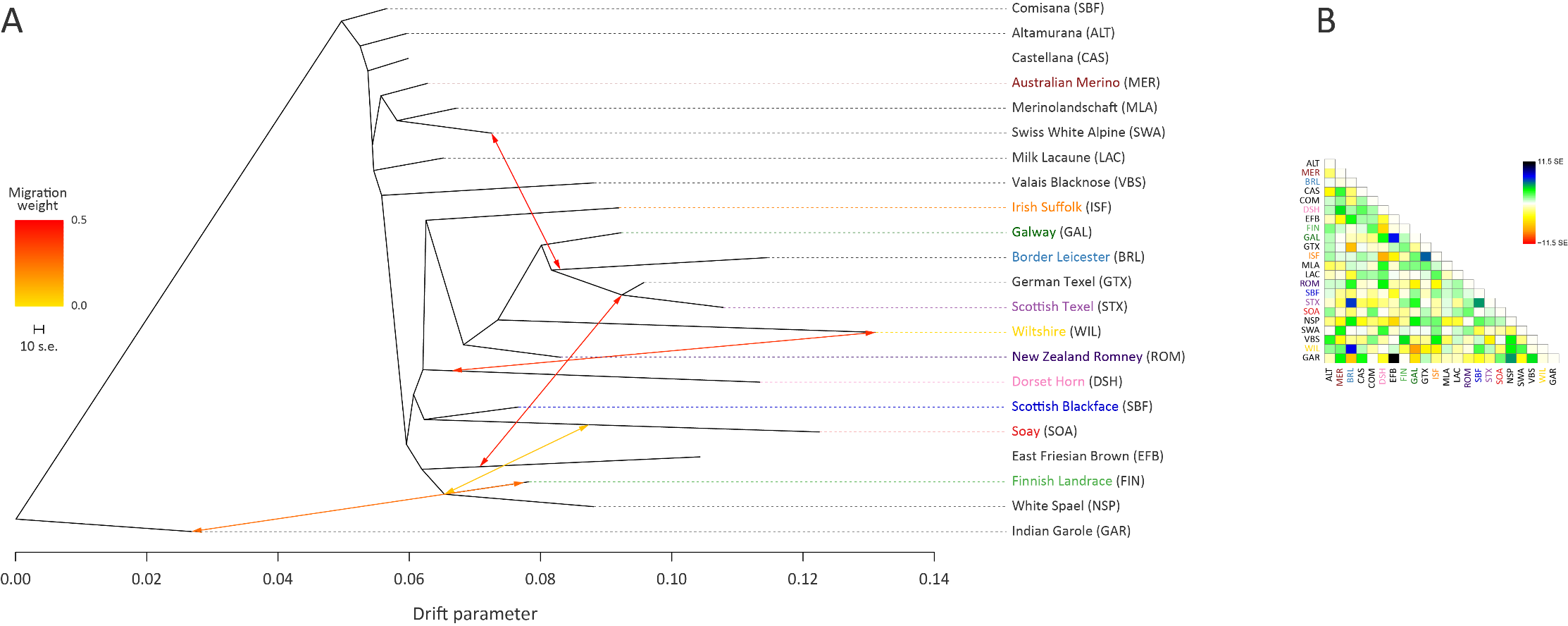

**Supplementary Figure 5.** Maximum likelihood (ML) phylogenetic tree network graph generated using 47,412 genome-wide SNPs with five migration edges showing the relationships among 22 sheep breeds (A), and the residuals indicating the fit of each pair of populations to the model (B). The arrows indicate gene flow events between the populations and the colours of the arrows indicate the relative weights of migration. Breed names in coloured text indicate breeds used in the other analyses.

#### 1.2. Supplementary Tables

**Supplementary Table 1:** Breed name, code, status (according to the FAO DAD-IS database), breed origin, observed heterozygosity (calculated using PLINK v1.07) and sample size for the core data set and the extended data set.

| **Breed name** | **Breed code** | **Breed origin** | **Breed**  **status** | **Observed heterozygosity** | ***n*** |
| --- | --- | --- | --- | --- | --- |
| Border Leicester | BRL | England | At risk | 0.291 | 48 |
| Dorset Horn | DSH | England | At risk | 0.316 | 21 |
| Finnish Landrace | FIN | Finland | Not at risk | 0.346 | 99 |
| Galway | GAL | Ireland | At risk | 0.336 | 49 |
| Irish Suffolk† | ISF | England | Not at risk | 0.314 | 55 |
| Australian Merino | MER | Spain / Unknown | Not at risk | 0.363 | 50 |
| New Zealand Romney | ROM | England | Not at risk | 0.347 | 24 |
| Scottish Blackface | SBF | Scotland | Not at risk | 0.361 | 56 |
| Scottish Texel | STX | Netherlands | Not at risk | 0.351 | 80 |
| Soay | SOA | Scotland | At risk | 0.267 | 110 |
| Wiltshire | WIL | England | At risk | 0.269 | 23 |
| **Core data set total** | | | | | **615** |
| Altamurana | ALT | Italy | At risk | - | 24 |
| Castellana | CAS | Spain | Not at risk | - | 23 |
| Comisana | COM | Sicily | Not at risk | - | 24 |
| East Friesian Brown | EFB | Germany | At risk | - | 39 |
| German Texel | GTX | Netherlands | Not at risk | - | 46 |
| Indian Garole | GAR | India | Unknown | - | 26 |
| Merinolandschaft | MLA | Germany | Not at risk | - | 24 |
| Milk Lacaune | LAC | France | Not at risk | - | 103 |
| White Spael | NSP | Norway | Not at risk | - | 32 |
| Swiss White Alpine | SWA | Switzerland | Not at risk | - | 24 |
| Valais Blacknose | VBS | Switzerland | Not at risk | - | 24 |
| **Extended data set total** | | | | | **1004** |

†One animal was removed as the evidence suggested that it had been mislabelled

**Supplementary Table 2:** Weighted estimates of *F*_ST_ calculated for each population pair using PLINK 1.9.

| **Breed** | **MER** | **BRL** | **DSH** | **FIN** | **GAL** | **ISF** | **ROM** | **SBF** | **STX** | **SOA** |
| --- | --- | --- | --- | --- | --- | --- | --- | --- | --- | --- |
| **BRL** | 0.174 |  |  |  |  |  |  |  |  |  |
| **DSH** | 0.160 | 0.264 |  |  |  |  |  |  |  |  |
| **FIN** | 0.084 | 0.182 | 0.177 |  |  |  |  |  |  |  |
| **GAL** | 0.118 | 0.132 | 0.202 | 0.129 |  |  |  |  |  |  |
| **ISF** | 0.118 | 0.209 | 0.209 | 0.130 | 0.152 |  |  |  |  |  |
| **ROM** | 0.096 | 0.167 | 0.183 | 0.110 | 0.110 | 0.136 |  |  |  |  |
| **SBF** | 0.080 | 0.179 | 0.169 | 0.091 | 0.121 | 0.121 | 0.100 |  |  |  |
| **STX** | 0.128 | 0.169 | 0.214 | 0.131 | 0.119 | 0.166 | 0.132 | 0.128 |  |  |
| **SOA** | 0.208 | 0.302 | 0.302 | 0.203 | 0.250 | 0.250 | 0.236 | 0.204 | 0.251 |  |
| **WIL** | 0.188 | 0.261 | 0.286 | 0.199 | 0.207 | 0.232 | 0.200 | 0.193 | 0.222 | 0.326 |

**Supplementary Table 3:** *P*-values for Kolmogorov-Smirnov tests for normality of the distribution of effective population size (*N_e_*), *P*-values for Shapiro-Wilk tests for normality of the distribution of genomic inbreeding coefficient (*F*) and *P*-values for Shapiro-Wilk tests for normality of the distribution of inbreeding coefficient calculated using ROH (*F*_ROH_) results for each breed calculated using R; *P*-values < 0.05 are highlighted.

| **Breed** | ***N_e_ P*-values** | ***F P*-values** | ***F*_ROH_ *P*-values** |
| --- | --- | --- | --- |
| BRL | 0.000 | 0.558 | 0.026 |
| DSH | 0.000 | 0.001 | 0.001 |
| FIN | 0.000 | 0.000 | 0.000 |
| GAL | 0.000 | 0.000 | 0.000 |
| ISF | 0.000 | 0.000 | 0.000 |
| MER | 0.000 | 0.000 | 0.000 |
| ROM | 0.000 | 0.000 | 0.000 |
| SBF | 0.000 | 0.077 | 0.001 |
| STX | 0.000 | 0.000 | 0.000 |
| SOA | 0.000 | 0.386 | 0.003 |
| WIL | 0.000 | 0.076 | 0.699 |

**Supplementary Table 4:** Historical effective population size (*N*_e_) estimates calculated using SNeP; breeds classified as “at risk” and are highlighted in orange with other breeds highlighted in green.

| **Gens**  **Ago** | **BRL** | **DSH** | **FIN** | **GAL** | **ISF** | **MER** | **ROM** | **SBF** | **STX** | **SOA** | **WIL** |
| --- | --- | --- | --- | --- | --- | --- | --- | --- | --- | --- | --- |
| **13** | 223 | 61 | 445 | 184 | 178 | 619 | 370 | 315 | 150 | 146 | 50 |
| **15** | 217 | 66 | 474 | 196 | 188 | 655 | 377 | 334 | 160 | 143 | 53 |
| **17** | 217 | 73 | 495 | 206 | 198 | 680 | 384 | 358 | 172 | 145 | 57 |
| **20** | 214 | 81 | 528 | 220 | 210 | 708 | 402 | 378 | 185 | 145 | 62 |
| **23** | 216 | 88 | 568 | 238 | 223 | 758 | 414 | 406 | 200 | 150 | 69 |
| **27** | 215 | 98 | 607 | 252 | 240 | 809 | 429 | 441 | 217 | 153 | 76 |
| **32** | 216 | 110 | 657 | 277 | 260 | 863 | 437 | 479 | 234 | 160 | 85 |
| **38** | 223 | 125 | 725 | 298 | 282 | 931 | 464 | 524 | 259 | 171 | 96 |
| **45** | 229 | 141 | 805 | 326 | 306 | 1021 | 488 | 579 | 285 | 182 | 109 |
| **54** | 242 | 163 | 897 | 355 | 342 | 1113 | 526 | 657 | 318 | 192 | 126 |
| **65** | 257 | 191 | 1005 | 398 | 389 | 1250 | 574 | 733 | 356 | 211 | 147 |
| **80** | 278 | 219 | 1154 | 443 | 439 | 1400 | 634 | 831 | 403 | 233 | 172 |
| **98** | 311 | 261 | 1308 | 510 | 515 | 1594 | 722 | 954 | 471 | 264 | 205 |
| **120** | 351 | 313 | 1484 | 590 | 596 | 1824 | 789 | 1097 | 545 | 298 | 238 |
| **150** | 400 | 376 | 1687 | 673 | 701 | 2049 | 911 | 1282 | 628 | 342 | 296 |
| **187** | 461 | 451 | 1957 | 774 | 821 | 2358 | 1066 | 1484 | 741 | 406 | 358 |
| **234** | 540 | 547 | 2190 | 934 | 985 | 2669 | 1224 | 1739 | 872 | 483 | 448 |
| **293** | 633 | 655 | 2444 | 1065 | 1175 | 2865 | 1417 | 1970 | 1012 | 562 | 531 |
| **366** | 747 | 797 | 2707 | 1245 | 1361 | 3044 | 1612 | 2223 | 1169 | 670 | 664 |
| **454** | 897 | 950 | 2987 | 1452 | 1599 | 3411 | 1834 | 2493 | 1383 | 796 | 775 |
| **552** | 1036 | 1135 | 3161 | 1682 | 1770 | 3545 | 2111 | 2684 | 1537 | 933 | 956 |
| **657** | 1146 | 1262 | 3357 | 1811 | 2043 | 3731 | 2303 | 2889 | 1715 | 1082 | 1080 |
| **761** | 1334 | 1423 | 3441 | 2069 | 2275 | 3783 | 2447 | 3080 | 1893 | 1224 | 1227 |
| **847** | 1392 | 1579 | 3696 | 2087 | 2317 | 4088 | 2612 | 3134 | 2000 | 1224 | 1371 |
| **914** | 1501 | 1639 | 3674 | 2307 | 2436 | 4343 | 2713 | 3419 | 2174 | 1413 | 1431 |
| **959** | 1515 | 1710 | 3593 | 2262 | 2478 | 4143 | 2515 | 3385 | 2029 | 1303 | 1529 |
| **983** | **-** | **-** | 3932 | 2427 | 2684 | 3926 | 2500 | 3197 | 2002 | - | - |

**Supplementary Table 5:** Bonferroni-corrected *P*-values for Wilcoxon rank sum pairwise-tests of breed differences in effective population size (*N_e_*) calculated using R after a significant Kruskal-Wallis test (*H* = 87.076, df = 10, *P-*value < 0.001); *P*-values < 0.05 are highlighted.

| **Breed** | **BRL** | **DSH** | **FIN** | **GAL** | **ISF** | **MER** | **ROM** | **SBF** | **STX** | **SOA** |
| --- | --- | --- | --- | --- | --- | --- | --- | --- | --- | --- |
| **DSH** | 1.000 |  |  |  |  |  |  |  |  |  |
| **FIN** | 0.001 | 0.000 |  |  |  |  |  |  |  |  |
| **GAL** | 1.000 | 0.916 | 0.067 |  |  |  |  |  |  |  |
| **ISF** | 1.000 | 0.827 | 0.127 | 1.000 |  |  |  |  |  |  |
| **MER** | 0.000 | 0.000 | 1.000 | 0.006 | 0.012 |  |  |  |  |  |
| **ROM** | 0.036 | 0.042 | 1.000 | 1.000 | 1.000 | 0.092 |  |  |  |  |
| **SBF** | 0.011 | 0.006 | 1.000 | 1.000 | 1.000 | 1.000 | 1.000 |  |  |  |
| **STX** | 1.000 | 1.000 | 0.036 | 1.000 | 1.000 | 0.002 | 1.000 | 0.531 |  |  |
| **SOA** | 1.000 | 1.000 | 0.000 | 0.516 | 0.602 | 0.000 | 0.015 | 0.003 | 1.000 |  |
| **WIL** | 1.000 | 1.000 | 0.000 | 0.230 | 0.181 | 0.000 | 0.007 | 0.003 | 0.490 | 1.000 |

**Supplementary Table 6:** Median, minimum and maximum, for each breed, for the genomic inbreeding coefficient (*F*) and the *F*_ROH_ inbreeding coefficient.

| **Breed** | **Median**  ***F*** | **Minimum**  ***F*** | **Maximum**  ***F*** | **Median**  ***F*_ROH_** | **Minimum**  ***F*_ROH_** | **Maximum**  ***F*_ROH_** |
| --- | --- | --- | --- | --- | --- | --- |
| BRL | 0.243 | 0.170 | 0.305 | 0.091 | 0.045 | 0.174 |
| DSH | 0.169 | 0.115 | 0.389 | 0.144 | 0.083 | 0.339 |
| FIN | 0.087 | 0.041 | 0.293 | 0.032 | 0.002 | 0.261 |
| GAL | 0.127 | 0.072 | 0.294 | 0.050 | 0.011 | 0.238 |
| ISF | 0.185 | -0.009 | 0.367 | 0.106 | 0.000 | 0.319 |
| MER | 0.045 | 0.012 | 0.233 | 0.026 | 0.006 | 0.200 |
| ROM | 0.086 | 0.056 | 0.211 | 0.041 | 0.015 | 0.151 |
| SBF | 0.060 | 0.022 | 0.128 | 0.029 | 0.005 | 0.080 |
| STX | 0.111 | -0.030 | 0.175 | 0.045 | 0.000 | 0.110 |
| SOA | 0.308 | 0.261 | 0.356 | 0.101 | 0.058 | 0.182 |
| WIL | 0.299 | 0.197 | 0.362 | 0.263 | 0.165 | 0.319 |

**Supplementary Table 7:** Bonferroni-corrected *P*-values for Wilcoxon rank sum pairwise-tests on breed differences in genomic inbreeding coefficient (*F*) calculated using R after a significant Kruskal-Wallis test (*H* = 477.33, df = 10, *P-*value < 0.001); *P*-values < 0.05 are highlighted.

| **Breed** | **BRL** | **DSH** | **FIN** | **GAL** | **ISF** | **MER** | **ROM** | **SBF** | **STX** | **SOA** |
| --- | --- | --- | --- | --- | --- | --- | --- | --- | --- | --- |
| **DSH** | 0.000 |  |  |  |  |  |  |  |  |  |
| **FIN** | 0.000 | 0.000 |  |  |  |  |  |  |  |  |
| **GAL** | 0.000 | 0.006 | 0.002 |  |  |  |  |  |  |  |
| **ISF** | 0.000 | 1.000 | 0.000 | 0.000 |  |  |  |  |  |  |
| **MER** | 0.000 | 0.000 | 1.000 | 0.000 | 0.000 |  |  |  |  |  |
| **ROM** | 0.000 | 0.000 | 1.000 | 0.074 | 0.000 | 0.000 |  |  |  |  |
| **SBF** | 0.000 | 0.000 | 0.000 | 0.000 | 0.000 | 1.000 | 0.000 |  |  |  |
| **STX** | 0.000 | 0.000 | 1.000 | 0.298 | 0.000 | 0.003 | 1.000 | 0.000 |  |  |
| **SOA** | 0.000 | 0.000 | 0.000 | 0.000 | 0.000 | 0.000 | 0.000 | 0.000 | 0.000 |  |
| **WIL** | 0.000 | 0.000 | 0.000 | 0.000 | 0.000 | 0.000 | 0.000 | 0.000 | 0.000 | 1.000 |

**Supplementary Table 8:** Bonferroni-corrected *P*-values for Wilcoxon rank sum pairwise-tests of breed differences in *F*_ROH_ inbreeding coefficient calculated using R after a significant Kruskal-Wallis test (*H* = 345.6, df = 10, *P-*value < 0.001); *P*-values < 0.05 are highlighted.

| **Breed** | **BRL** | **DSH** | **FIN** | **GAL** | **ISF** | **MER** | **ROM** | **SBF** | **STX** | **SOA** |
| --- | --- | --- | --- | --- | --- | --- | --- | --- | --- | --- |
| **DSH** | 0.000 |  |  |  |  |  |  |  |  |  |
| **FIN** | 0.000 | 0.000 |  |  |  |  |  |  |  |  |
| **GAL** | 0.000 | 0.000 | 0.155 |  |  |  |  |  |  |  |
| **ISF** | 0.728 | 0.112 | 0.000 | 0.000 |  |  |  |  |  |  |
| **MER** | 0.000 | 0.000 | 1.000 | 0.018 | 0.000 |  |  |  |  |  |
| **ROM** | 0.000 | 0.000 | 1.000 | 1.000 | 0.000 | 1.000 |  |  |  |  |
| **SBF** | 0.000 | 0.000 | 1.000 | 0.001 | 0.000 | 1.000 | 0.389 |  |  |  |
| **STX** | 0.000 | 0.000 | 1.000 | 1.000 | 0.000 | 1.000 | 1.000 | 1.000 |  |  |
| **SOA** | 1.000 | 0.000 | 0.000 | 0.000 | 1.000 | 0.000 | 0.000 | 0.000 | 0.000 |  |
| **WIL** | 0.000 | 0.000 | 0.000 | 0.000 | 0.000 | 0.000 | 0.000 | 0.000 | 0.000 | 0.000 |

**Supplementary Table 9:** The location of region for the clusters of SNPs with the top 0.1% CSS, number of SNPs and maximum CSS *P*-value in the clusters, number of genes in the 1 Mb flanking the clusters upstream and downstream and IDs of Galway sheep within the flanking region included in a ROH.

| **Cluster Region**  **(Mb range)** | **No. of SNPs in cluster** | **Maximum CSS in cluster** | **No of genes in flanking 1Mb** | **Galway sheep with ROH in flanking 1 Mb** |
| --- | --- | --- | --- | --- |
| OAR1: 42.29 -43.08 | 14 | 3.917 | 23 | GAL2, GAL15, GAL26, GAL49, GAL50 |
| OAR1: 102.45 -103.2 | 14 | 3.785 | 104 | GAL11, GAL13 GAL45 |
| OAR3: 40.64 -40.68 | 2 | 3.367 | 13 | GAL4, GAL12, GAL 19, GAL27, GAL35, GAL44 |
| OAR8: 49.15 -49.85 | 16 | 4.027 | 30 | GAL18, GAL30, GAL35, GAL36, GAL38, GAL45, GAL50 |
| OAR8: 56.82 -56.88 | 2 | 3.390 | 26 | GAL15, GAL18, GAL26, GAL30, GAL45, GAL46 |

**Supplementary Table 10:** The location of selection peak cluster regions, Ensembl ID, HGNC symbol and description of the candidate genes under selection in the Galway breed.

| Cluster region (Mb range) | Ensembl ID | Gene symbol | Description |
| --- | --- | --- | --- |
| OAR1:42.29-43.08 | ENSOARG00000010091 | *-* | Uncharacterized protein |
| OAR1:42.29-43.08 | ENSOARG00000010122 | *DIRAS3* | DIRAS family GTPase 3 |
| OAR1:42.29-43.08 | ENSOARG00000010521 | *PDE4B* | phosphodiesterase 4B |
| OAR1:42.29-43.08 | ENSOARG00000010617 | *SGIP1* | SH3 domain GRB2 like endophilin interacting protein 1 |
| OAR1:42.29-43.08 | ENSOARG00000010702 | *TCTEX1D1* | Tctex1 domain containing 1 |
| OAR1:42.29-43.08 | ENSOARG00000010717 | *INSL5* | insulin like 5 |
| OAR1:42.29-43.08 | ENSOARG00000010792 | *WDR78* | WD repeat domain 78 |
| OAR1:42.29-43.08 | ENSOARG00000010903 | *MIER1* | MIER1 transcriptional regulator |
| OAR1:42.29-43.08 | ENSOARG00000010986 | *SLC35D1* | solute carrier family 35 member D1 |
| OAR1:42.29-43.08 | ENSOARG00000011004 | *C1orf141* | chromosome 1 open reading frame 141 |
| OAR1:42.29-43.08 | ENSOARG00000011083 | *IL23R* | interleukin 23 receptor |
| OAR1:42.29-43.08 | ENSOARG00000011173 | *IL12RB2* | interleukin 12 receptor subunit beta 2 |
| OAR1:42.29-43.08 | ENSOARG00000011282 | *SERBP1* | SERPINE1 mRNA binding protein 1 |
| OAR1:42.29-43.08 | ENSOARG00000011356 | *GADD45A* | growth arrest and DNA damage inducible alpha |
| OAR1:42.29-43.08 | ENSOARG00000011369 | *GNG12* | G protein subunit gamma 12 |
| OAR1:42.29-43.08 | ENSOARG00000011383 | *WLS* | wntless Wnt ligand secretion mediator |
| OAR1:42.29-43.08 | ENSOARG00000011470 | *RPE65* | retinal pigment epithelium-specific protein 65kDa |
| OAR1:42.29-43.08 | ENSOARG00000011569 | *DEPDC1* | DEP domain containing 1 |
| OAR1:42.29-43.08 | ENSOARG00000021718 | *-* | Small nucleolar RNA SNORA70 |
| OAR1:42.29-43.08 | ENSOARG00000023397 | *-* | U6 spliceosomal RNA |
| OAR1:42.29-43.08 | ENSOARG00000024412 | *-* | - |
| OAR1:42.29-43.08 | ENSOARG00000025521 | *-* | - |
| OAR1:42.29-43.08 | ENSOARG00000025522 | *-* | - |
| OAR1:102.45-103.2 | ENSOARG00000000259 | *C1orf68* | chromosome 1 open reading frame 68 |
| OAR1:102.45-103.2 | ENSOARG00000000272 | *-* | Uncharacterized protein |
| OAR1:102.45-103.2 | ENSOARG00000000287 | *KPRP* | keratinocyte proline rich protein |
| OAR1:102.45-103.2 | ENSOARG00000000297 | *-* | Uncharacterized protein |
| OAR1:102.45-103.2 | ENSOARG00000000314 | *-* | Uncharacterized protein |
| OAR1:102.45-103.2 | ENSOARG00000000330 | *-* | - |
| OAR1:102.45-103.2 | ENSOARG00000000342 | *SPRR3* | small proline rich protein 3 |
| OAR1:102.45-103.2 | ENSOARG00000000351 | *-* | Uncharacterized protein |
| OAR1:102.45-103.2 | ENSOARG00000000368 | *-* | - |
| OAR1:102.45-103.2 | ENSOARG00000000377 | *-* | Uncharacterized protein |
| OAR1:102.45-103.2 | ENSOARG00000000389 | *PGLYRP3* | peptidoglycan recognition protein 3 |
| OAR1:102.45-103.2 | ENSOARG00000000406 | *PGLYRP4* | peptidoglycan recognition protein 4 |
| OAR1:102.45-103.2 | ENSOARG00000000417 | *S100A9* | S100 calcium binding protein A9 |
| OAR1:102.45-103.2 | ENSOARG00000000432 | *S100A12* | S100 calcium binding protein A12 |
| OAR1:102.45-103.2 | ENSOARG00000000450 | *S100A8* | S100 calcium binding protein A8 |
| OAR1:102.45-103.2 | ENSOARG00000000463 | *-* | Uncharacterized protein |
| OAR1:102.45-103.2 | ENSOARG00000000477 | *-* | Uncharacterized protein |
| OAR1:102.45-103.2 | ENSOARG00000000593 | *-* | Protein S100 |
| OAR1:102.45-103.2 | ENSOARG00000000610 | *S100A6* | S100 calcium binding protein A6 |
| OAR1:102.45-103.2 | ENSOARG00000000623 | *S100A5* | S100 calcium binding protein A5 |
| OAR1:102.45-103.2 | ENSOARG00000000641 | *S100A4* | S100 calcium binding protein A4 |
| OAR1:102.45-103.2 | ENSOARG00000000657 | *S100A3* | S100 calcium binding protein A3 |
| OAR1:102.45-103.2 | ENSOARG00000000674 | *S100A2* | S100 calcium binding protein A2 |
| OAR1:102.45-103.2 | ENSOARG00000000691 | *S100A16* | S100 calcium binding protein A16 |
| OAR1:102.45-103.2 | ENSOARG00000000707 | *S100A1* | S100 calcium binding protein A1 |
| OAR1:102.45-103.2 | ENSOARG00000000735 | *CHTOP* | chromatin target of PRMT1 |
| OAR1:102.45-103.2 | ENSOARG00000000758 | *SNAPIN* | SNAP associated protein |
| OAR1:102.45-103.2 | ENSOARG00000000787 | *ILF2* | interleukin enhancer binding factor 2 |
| OAR1:102.45-103.2 | ENSOARG00000000858 | *NPR1* | natriuretic peptide receptor 1 |
| OAR1:102.45-103.2 | ENSOARG00000001095 | *INTS3* | integrator complex subunit 3 |
| OAR1:102.45-103.2 | ENSOARG00000001265 | *SLC27A3* | solute carrier family 27 member 3 |
| OAR1:102.45-103.2 | ENSOARG00000001449 | *-* | Uncharacterized protein |
| OAR1:102.45-103.2 | ENSOARG00000001486 | *GATAD2B* | GATA zinc finger domain containing 2B |
| OAR1:102.45-103.2 | ENSOARG00000001584 | *DENND4B* | DENN domain containing 4B |
| OAR1:102.45-103.2 | ENSOARG00000001610 | *-* | Uncharacterized protein |
| OAR1:102.45-103.2 | ENSOARG00000001630 | *SLC39A1* | solute carrier family 39 member 1 |
| OAR1:102.45-103.2 | ENSOARG00000001743 | *CREB3L4* | cAMP responsive element binding protein 3 like 4 |
| OAR1:102.45-103.2 | ENSOARG00000001762 | *JTB* | jumping translocation breakpoint |
| OAR1:102.45-103.2 | ENSOARG00000001777 | *RAB13* | RAB13, member RAS oncogene family |
| OAR1:102.45-103.2 | ENSOARG00000001813 | *NUP210L* | nucleoporin 210 like |
| OAR1:102.45-103.2 | ENSOARG00000001845 | *TPM3* | tropomyosin 3 |
| OAR1:102.45-103.2 | ENSOARG00000001925 | *C1orf189* | chromosome 1 open reading frame 189 |
| OAR1:102.45-103.2 | ENSOARG00000002028 | *-* | Uncharacterized protein |
| OAR1:102.45-103.2 | ENSOARG00000002170 | *UBAP2L* | ubiquitin associated protein 2 like |
| OAR1:102.45-103.2 | ENSOARG00000002271 | *HAX1* | HCLS1 associated protein X-1 |
| OAR1:102.45-103.2 | ENSOARG00000002349 | *AQP10* | aquaporin 10 |
| OAR1:102.45-103.2 | ENSOARG00000002448 | *ATP8B2* | ATPase phospholipid transporting 8B2 |
| OAR1:102.45-103.2 | ENSOARG00000002562 | *IL6R* | interleukin 6 receptor |
| OAR1:102.45-103.2 | ENSOARG00000002674 | *SHE* | Src homology 2 domain containing E |
| OAR1:102.45-103.2 | ENSOARG00000002772 | *TDRD10* | tudor domain containing 10 |
| OAR1:102.45-103.2 | ENSOARG00000002800 | *UBE2Q1* | ubiquitin conjugating enzyme E2 Q1 |
| OAR1:102.45-103.2 | ENSOARG00000002921 | *CHRNB2* | cholinergic receptor nicotinic beta 2 subunit |
| OAR1:102.45-103.2 | ENSOARG00000002952 | *ADAR* | adenosine deaminase, RNA specific |
| OAR1:102.45-103.2 | ENSOARG00000002973 | *KCNN3* | potassium calcium-activated channel subfamily N member 3 |
| OAR1:102.45-103.2 | ENSOARG00000003056 | *PMVK* | phosphomevalonate kinase |
| OAR1:102.45-103.2 | ENSOARG00000003126 | *PBXIP1* | PBX homeobox interacting protein 1 |
| OAR1:102.45-103.2 | ENSOARG00000003199 | *SHC1* | SHC adaptor protein 1 |
| OAR1:102.45-103.2 | ENSOARG00000003292 | *CKS1B* | CDC28 protein kinase regulatory subunit 1B |
| OAR1:102.45-103.2 | ENSOARG00000003310 | *FLAD1* | flavin adenine dinucleotide synthetase 1 |
| OAR1:102.45-103.2 | ENSOARG00000003333 | *ZBTB7B* | zinc finger and BTB domain containing 7B |
| OAR1:102.45-103.2 | ENSOARG00000003355 | *DCST2* | DC-STAMP domain containing 2 |
| OAR1:102.45-103.2 | ENSOARG00000003448 | *DCST1* | DC-STAMP domain containing 1 |
| OAR1:102.45-103.2 | ENSOARG00000003543 | *ADAM15* | ADAM metallopeptidase domain 15 |
| OAR1:102.45-103.2 | ENSOARG00000003565 | *EFNA4* | ephrin A4 |
| OAR1:102.45-103.2 | ENSOARG00000003579 | *-* | Uncharacterized protein |
| OAR1:102.45-103.2 | ENSOARG00000003647 | *EFNA1* | ephrin A1 |
| OAR1:102.45-103.2 | ENSOARG00000003734 | *SLC50A1* | solute carrier family 50 member 1 |
| OAR1:102.45-103.2 | ENSOARG00000003747 | *KRTCAP2* | keratinocyte associated protein 2 |
| OAR1:102.45-103.2 | ENSOARG00000003769 | *TRIM46* | tripartite motif containing 46 |
| OAR1:102.45-103.2 | ENSOARG00000003788 | *MUC1* | mucin 1, cell surface associated |
| OAR1:102.45-103.2 | ENSOARG00000003894 | *THBS3* | thrombospondin 3 |
| OAR1:102.45-103.2 | ENSOARG00000003912 | *-* | Uncharacterized protein |
| OAR1:102.45-103.2 | ENSOARG00000003937 | *GBA* | glucosylceramidase beta |
| OAR1:102.45-103.2 | ENSOARG00000003954 | *FAM189B* | family with sequence similarity 189 member B |
| OAR1:102.45-103.2 | ENSOARG00000004032 | *SCAMP3* | secretory carrier membrane protein 3 |
| OAR1:102.45-103.2 | ENSOARG00000004102 | *CLK2* | CDC like kinase 2 |
| OAR1:102.45-103.2 | ENSOARG00000004200 | *HCN3* | hyperpolarization activated cyclic nucleotide gated potassium channel 3 |
| OAR1:102.45-103.2 | ENSOARG00000004229 | *PKLR* | pyruvate kinase, liver and RBC |
| OAR1:102.45-103.2 | ENSOARG00000004341 | *-* | - |
| OAR1:102.45-103.2 | ENSOARG00000004448 | *ASH1L* | ASH1 like histone lysine methyltransferase |
| OAR1:102.45-103.2 | ENSOARG00000006463 | *-* | Uncharacterized protein |
| OAR1:102.45-103.2 | ENSOARG00000006502 | *-* | Uncharacterized protein |
| OAR1:102.45-103.2 | ENSOARG00000006529 | *-* | - |
| OAR1:102.45-103.2 | ENSOARG00000006552 | *-* | Uncharacterized protein |
| OAR1:102.45-103.2 | ENSOARG00000006578 | *PRR9* | proline rich 9 |
| OAR1:102.45-103.2 | ENSOARG00000006605 | *-* | Uncharacterized protein |
| OAR1:102.45-103.2 | ENSOARG00000006634 | *LENEP* | lens epithelial protein |
| OAR1:102.45-103.2 | ENSOARG00000006665 | *DPM3* | dolichyl-phosphate mannosyltransferase subunit 3 |
| OAR1:102.45-103.2 | ENSOARG00000022891 | *-* | U6 spliceosomal RNA |
| OAR1:102.45-103.2 | ENSOARG00000023246 | *-* | Small nucleolar RNA SNORA51 |
| OAR1:102.45-103.2 | ENSOARG00000023446 | *-* | Small nucleolar RNA SNORA58 |
| OAR1:102.45-103.2 | ENSOARG00000024783 | *-* | U6 spliceosomal RNA |
| OAR1:102.45-103.2 | ENSOARG00000024820 | *-* | - |
| OAR1:102.45-103.2 | ENSOARG00000025019 | *-* | Small nucleolar RNA SNORA58 |
| OAR1:102.45-103.2 | ENSOARG00000025190 | *-* | small proline-rich protein type II |
| OAR1:102.45-103.2 | ENSOARG00000025191 | *-* | Uncharacterized protein |
| OAR1:102.45-103.2 | ENSOARG00000025193 | *-* | Uncharacterized protein |
| OAR1:102.45-103.2 | ENSOARG00000025195 | *-* | Uncharacterized protein |
| OAR1:102.45-103.2 | ENSOARG00000025551 | *-* | - |
| OAR1:102.45-103.2 | ENSOARG00000025552 | *-* | - |
| OAR1:102.45-103.2 | ENSOARG00000025553 | *-* | - |
| OAR1:102.45-103.2 | ENSOARG00000025554 | *-* | - |
| OAR1:102.45-103.2 | ENSOARG00000025555 | *-* | - |
| OAR1:102.45-103.2 | ENSOARG00000025556 | *-* | - |
| OAR3:40.64-40.68 | ENSOARG00000019897 | *APLF* | aprataxin and PNKP like factor |
| OAR3:40.64-40.68 | ENSOARG00000019899 | *FBXO48* | F-box protein 48 |
| OAR3:40.64-40.68 | ENSOARG00000019902 | *PLEK* | pleckstrin |
| OAR3:40.64-40.68 | ENSOARG00000019907 | *CNRIP1* | cannabinoid receptor interacting protein 1 |
| OAR3:40.64-40.68 | ENSOARG00000019910 | *-* | Uncharacterized protein |
| OAR3:40.64-40.68 | ENSOARG00000019914 | *PNO1* | partner of NOB1 homolog |
| OAR3:40.64-40.68 | ENSOARG00000019929 | *-* | Uncharacterized protein |
| OAR3:40.64-40.68 | ENSOARG00000019931 | *C1D* | C1D nuclear receptor corepressor |
| OAR3:40.64-40.68 | ENSOARG00000019948 | *ETAA1* | Ewing tumor associated antigen 1 |
| OAR3:40.64-40.68 | ENSOARG00000019962 | *MEIS1* | Meis homeobox 1 |
| OAR3:40.64-40.68 | ENSOARG00000024135 | *-* | - |
| OAR3:40.64-40.68 | ENSOARG00000025922 | *-* | - |
| OAR3:40.64-40.68 | ENSOARG00000025970 | *-* | - |
| OAR8:49.15-49.85 | ENSOARG00000012863 | *SRSF12* | serine and arginine rich splicing factor 12 |
| OAR8:49.15-49.85 | ENSOARG00000012871 | *PNRC1* | proline rich nuclear receptor coactivator 1 |
| OAR8:49.15-49.85 | ENSOARG00000012911 | *RNGTT* | RNA guanylyltransferase and 5'-phosphatase |
| OAR8:49.15-49.85 | ENSOARG00000012955 | *SPACA1* | sperm acrosome associated 1 |
| OAR8:49.15-49.85 | ENSOARG00000012963 | *-* | Uncharacterized protein |
| OAR8:49.15-49.85 | ENSOARG00000012967 | *AKIRIN2* | akirin 2 |
| OAR8:49.15-49.85 | ENSOARG00000012992 | *ORC3* | origin recognition complex subunit 3 |
| OAR8:49.15-49.85 | ENSOARG00000013003 | *RARS2* | arginyl-tRNA synthetase 2, mitochondrial |
| OAR8:49.15-49.85 | ENSOARG00000013049 | *SLC35A1* | solute carrier family 35 member A1 |
| OAR8:49.15-49.85 | ENSOARG00000013071 | *-* | Uncharacterized protein |
| OAR8:49.15-49.85 | ENSOARG00000013104 | *C6orf163* | chromosome 6 open reading frame 163 |
| OAR8:49.15-49.85 | ENSOARG00000013115 | *-* | Uncharacterized protein |
| OAR8:49.15-49.85 | ENSOARG00000013132 | *ZNF292* | zinc finger protein 292 |
| OAR8:49.15-49.85 | ENSOARG00000013153 | *CGA* | glycoprotein hormones, alpha polypeptide |
| OAR8:49.15-49.85 | ENSOARG00000013156 | *-* | Uncharacterized protein |
| OAR8:49.15-49.85 | ENSOARG00000019995 | *CNR1* | cannabinoid receptor 1 (brain) |
| OAR8:49.15-49.85 | ENSOARG00000020002 | *-* | Uncharacterized protein |
| OAR8:49.15-49.85 | ENSOARG00000020007 | *HTR1E* | 5-hydroxytryptamine receptor 1E |
| OAR8:49.15-49.85 | ENSOARG00000020008 | *-* | Uncharacterized protein |
| OAR8:49.15-49.85 | ENSOARG00000020012 | *-* | Uncharacterized protein |
| OAR8:49.15-49.85 | ENSOARG00000021301 | *-* | Small nucleolar RNA SNORA70 |
| OAR8:49.15-49.85 | ENSOARG00000021683 | *-* | U6 spliceosomal RNA |
| OAR8:49.15-49.85 | ENSOARG00000023346 | *-* | 5S ribosomal RNA |
| OAR8:49.15-49.85 | ENSOARG00000024253 | *-* | - |
| OAR8:49.15-49.85 | ENSOARG00000024586 | *-* | Small nucleolar RNA SNORD22 |
| OAR8:49.15-49.85 | ENSOARG00000024732 | *-* | U6 spliceosomal RNA |
| OAR8:49.15-49.85 | ENSOARG00000027025 | *-* | - |
| OAR8:49.15-49.85 | ENSOARG00000027026 | *-* | - |
| OAR8:49.15-49.85 | ENSOARG00000027027 | *-* | - |
| OAR8:49.15-49.85 | ENSOARG00000027028 | *-* | - |
